# ULK1/2 Inhibitors that Degrade ATG13 Effectively Target KRAS-Mutant Cancers

**DOI:** 10.1101/2025.10.14.682402

**Authors:** Patrick M. Hagan, Huiyu Ren, Sonja N. Brun, Fabiana I. A. Layng, Dhanya R. Panickar, Lester J. Lambert, Maria Celeridad, Ranajit Das, Li Ling, Yijuan Zhang, Allison S. Limpert, Douglas J. Sheffler, Cosimo Commisso, Reuben J. Shaw, Nicholas D. P. Cosford

## Abstract

KRAS mutations drive tumorigenesis in multiple cancer types, including lung and pancreatic cancer. Autophagy is a cell survival pathway that supports tumor growth under metabolic stress and has been proposed to be a potential therapeutic avenue specifically in KRAS mutant cancers. The Unc-51-like ATG-activating kinases 1 and 2 (ULK) initiate the earliest regulated steps of autophagy and are the only protein kinases in the canonical autophagy pathway, thus making them attractive therapeutic targets for KRAS mutant tumors. We show here that genetic depletion of ULK1 or ATG13, core components of the ULK1 complex, in KRAS mutant lung and pancreatic cancer cell lines results in growth inhibition. Previously, we developed small molecule ULK1 inhibitors that not only inhibit ULK kinase activity but also induced the degradation of other core members of the ULK complex including ATG101 and ATG13. Therefore, we developed a high-throughput screening (HTS) assay in which ATG13 was HiBiT-tagged in KRAS mutant lung cancer cells to evaluate ULK inhibitors for ATG13 degradation. Using this approach, we discovered a lead ULK inhibitor, **SBP-1750**, that potently inhibited ULK activity, promoted robust ATG13 degradation, impaired ATG, and induced KRAS mutant cancer cell death. Studies in a KRAS-mutant orthotopic syngeneic pancreatic cancer model show that oral treatment with **SBP-1750** significantly reduced tumor growth. Pharmacokinetic analysis of **SBP-1750** indicates favorable drug exposure and pharmacodynamic analysis confirms ATG13 degradation in vivo, mirroring in vitro results. Finally, immunohistochemical staining of orthotopic pancreatic tumors reveals a significant increase in CD4⁺ and CD8⁺ T cell infiltration upon treatment, suggesting that **SBP-1750** enhances anti-tumor immunity. These findings support further development of **SBP-1750** as a novel ATG-targeting cancer therapy.

## Introduction

Autophagy upregulation is implicated in several contexts and diseases, including cancer ^1,2^. In KRAS mutant cancers, autophagy plays a critical role in sustaining tumor growth by maintaining metabolic homeostasis under nutrient and oxygen-deprived conditions. KRAS-driven tumors exhibit heightened metabolic demands, and autophagy helps meet these demands by recycling proteins, lipids, and other cellular components to replenish key metabolites like acetyl-CoA, which are often depleted in these tumors ^3–5^. For example, in genetically engineered mouse models (GEMMs) of KRAS-driven cancers, genetic deletion of the essential autophagy gene ATG7 inhibits tumor growth. Furthermore, conditional deletion of ATG7 in pre-established, KRAS-dependent tumors demonstrate that advanced tumors have an even stronger dependence on ATG, emphasizing its critical role in tumor maintenance ^6^. This suggests that autophagy is a metabolic vulnerability in KRAS mutant cancers and that pharmacologically targeting autophagy could represent an effective therapeutic strategy ^7, 8^.

Despite the therapeutic potential of autophagy inhibition, current clinical options are limited. Chloroquine (CQ) and hydroxychloroquine (HCQ), FDA-approved antimalarial drugs, are being evaluated in the clinic as autophagy inhibitors due to their effects on lysosomal activity. However, CQ and HCQ are not specific for autophagy and therefore have other associated toxicities including retinopathy and cardiotoxicity ^9^. The lack of potent and specific small-molecule ATG-selective inhibitors has hindered clinical translation, leaving a substantial unmet need in the treatment of KRAS mutant cancers, which account for a significant portion of cancer-related mortality¹¹. Developing novel, highly potent autophagy inhibitors capable of targeting key autophagy machinery could provide a promising therapeutic approach for KRAS mutant cancers.

Autophagy is a multistep process starting with the activation of a pre-initiation complex, consisting of autophagy related protein 13 (ATG13), unc-51-like autophagy activating kinase 1 (ULK1) or its homologue ULK2, Atg101 and focal adhesion kinase (FAK) family-interacting protein (FIP200) ^10, 11^. ULK1 and ULK2 are human serine/threonine kinases that regulate the initiation of autophagy by phosphorylating ATG13, FIP200, and additional downstream effectors. Their activation marks the first committed step of ATG, coordinating recruitment of membranes and assembly of the autophagy machinery that leads to autophagosome formation ^12^. ATG13 binds ULK1 and FIP200 at its C terminus, between residues 347 and 480, and ATG101 closer to its N terminus, between residues 112 and 220 ^13–15^. ATG13 binds with the C terminal early autophagy tethering (EAT) domain of ULK1 to recruit the initiation complex to sites of phagophore initiation ^16, 17^. This supramolecular assembly of the autophagy pre-initiation complex by ATG13 is essential for autophagy initiation and thus tumor cell survival ^11, 18, 19^. CRISPR knockout or siRNA knockdown of ATG13 inhibits ATG, demonstrated by reduced LC3-I conversion to LC3-II ^14, 20^. In fact, knockout of ATG13 in NSCLC H292 cells and other ATG-dependent cancers had a more dramatic effect on cell growth than knockout of the essential gene proliferating cell nuclear antigen (PCNA) ^21^. These prior studies describing the essential roles of ATG13 in autophagy and experiments showing the effects of loss of ATG13 in cancer cells support ATG13 degradation as a viable treatment to kill tumor cells.

In our previous work, we observed that ULK kinase inhibitors caused marked reduction of ATG13 protein levels⁵. Targeted protein degradation has emerged as a prominent modality in modern drug discovery ^22–24^. Proteolysis-targeting chimeras (PROTACs) and molecular glues, which induce selective degradation of proteins of interest via the ubiquitin–proteasome system (UPS), are now advancing through clinical evaluation ^25–28^. These compounds cause a sustained and drug driven elimination of the protein of interest potentially improving selectivity, reducing dose burden, and overcoming resistance tied to active-site occupancy ^24, 29^. Investigating whether this combination of ULK inhibition and ATG13 degradation is key to the efficacy seen with our ULK inhibitors provoked us to focus compound design and SAR studies on promoting ATG13 degradation. To identify compounds that induce ATG13 protein loss and therefore inhibit autophagy in a high-throughput manner, we inserted a small bioluminescent HiBiT tag onto the C-terminus of ATG13 and developed a high-throughput ATG13 degradation assay. Using this approach, we identified a novel lead compound **SBP-1750,** which we detail here with pharmacodynamic (PD) and pharmacokinetic (PK) analysis and therapeutic potential in vivo in an immunocompetent KRAS G12D mutant pancreatic cancer orthotopic model.

## RESULTS

### Structure-Activity Relationship (SAR) Studies

We previously reported the design and synthesis of **SBP-7455** ^30^, a compound developed to enhance ULK target engagement using the co-crystal structure of the ULK2 kinase domain bound to the chemical probe **SBI-0206965**, our earlier generation ULK1 chemical probe ^31^. **SBP-7455** exhibited improved binding affinity for ULK and showed enhanced biochemical and cellular potency compared to **SBI-0206965**. SAR analysis of **SBI-0206955** and **SBP-7455** revealed that replacing the ether group with an amino group at the C-4 position of the pyrimidine scaffold significantly enhanced ULK inhibitory activity. Based on these findings, further structure-based rational design and lead optimization guided by the co-crystal structure of the ULK2 kinase domain bound to the chemical probe **SBI-0206965** ^31^, along with in silico modeling led to the discovery of **SBP-7501** and **SBP-5147** (**Supplemental Figure 1**) ^30–32^. Both compounds demonstrated improved biochemical potency and increased intracellular binding to ULK, compared to our previously published chemical probes **SBI-0206965** ^31^ and **SBP-7455** ^30^ (**Supplemental Figure 1 and Table 1**). Both **SBP-5147** and **SBP-7501** demonstrate strong cytotoxic effects against NSCLC cells. Moreover, **SBP-5147** effectively inhibited autophagy in vitro and enhanced the expression of major histocompatibility complex (MHC) class I in NSCLC cells (**Table 1**) ^32^. As part of our ongoing efforts to identify novel ULK inhibitors, herein we report further SAR studies of our first-generation chemical probe **SBI-0206965**. In our current study, we designed and synthesized a focused library of compounds by replacing the C-4 ether substituent of **SBI-0206965** with the corresponding 2-amino N-methylbenzamide moiety, along with the introduction of various N^2^-amino groups that had previously shown good in vitro potency in in-house assays.

**Table 1:**
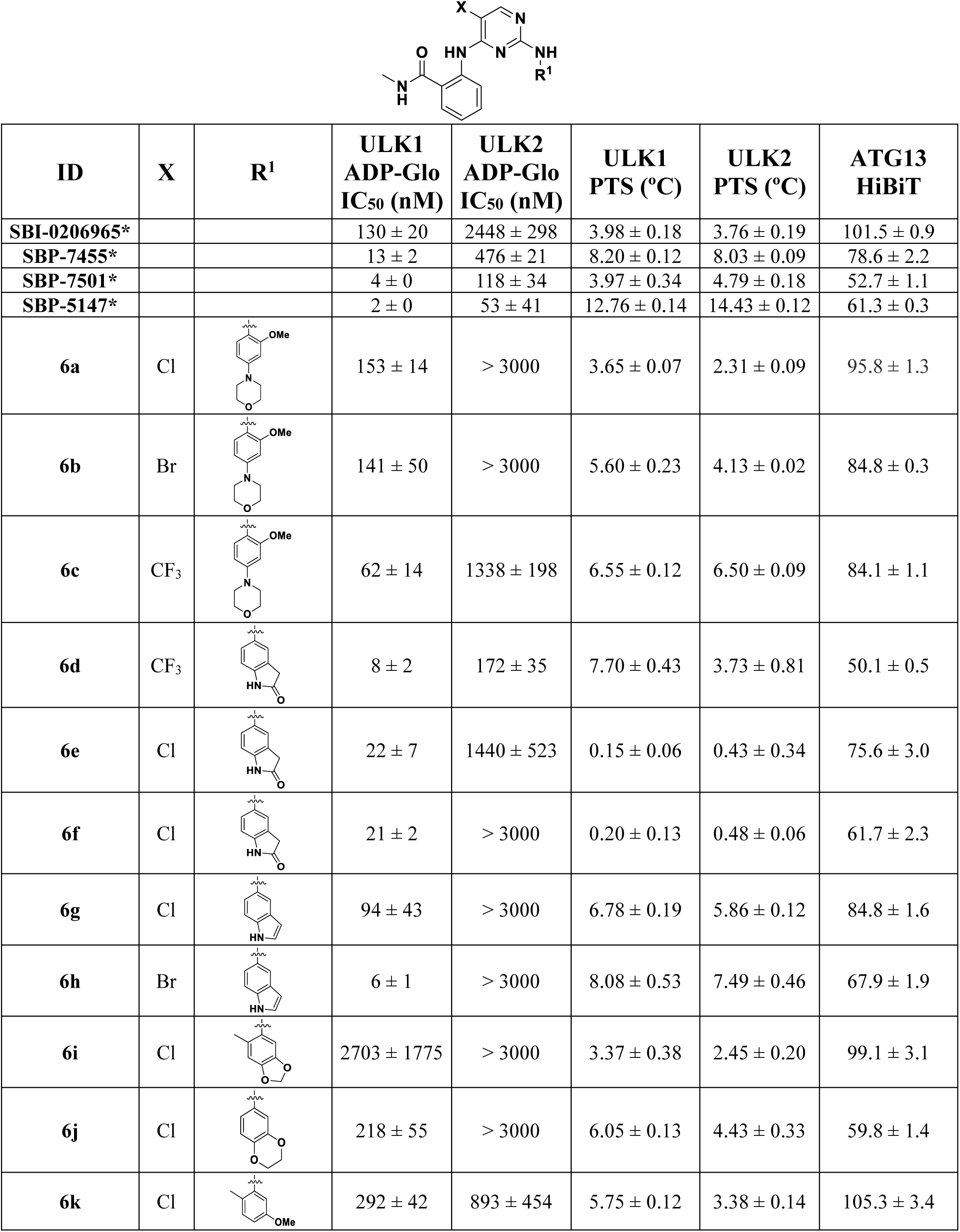
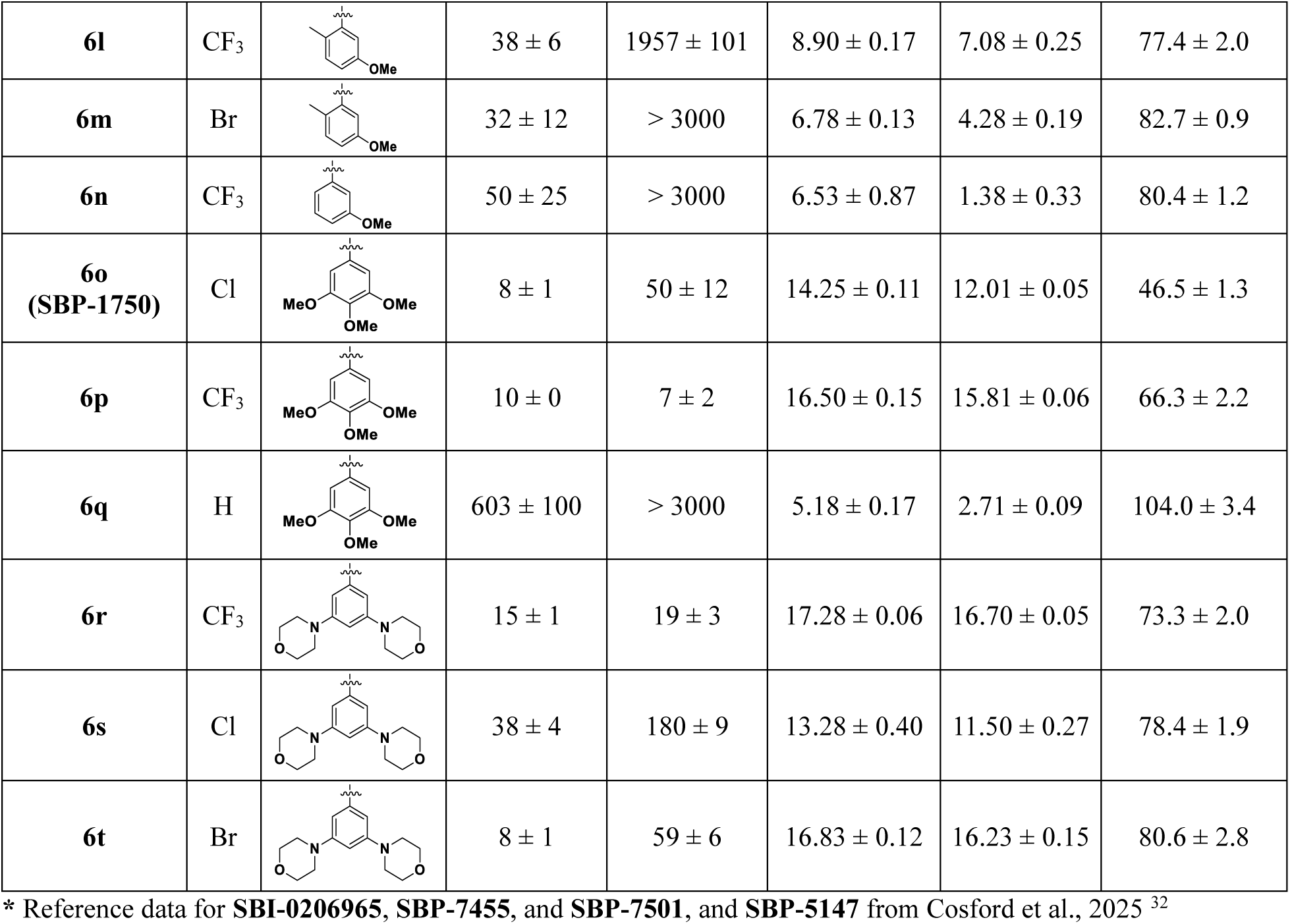
The structure of designed compounds, ULK, PTS and ATG13 HiBiT (% ATG13 Expression) data.

| ID | X | R<sup>1</sup> | ULK1<br>ADP-Glo<br>IC<sub>50</sub> (nM) | ULK2<br>ADP-Glo<br>IC<sub>50</sub> (nM) | ULK1<br>PTS (°C) | ULK2<br>PTS (°C) | ATG13<br>HiBiT |
| --- | --- | --- | --- | --- | --- | --- | --- |
| **SBI-0206965\*** |  |  | 130 ± 20 | 2448 ± 298 | 3.98 ± 0.18 | 3.76 ± 0.19 | 101.5 ± 0.9 |
| **SBP-7455\*** |  |  | 13 ± 2 | 476 ± 21 | 8.20 ± 0.12 | 8.03 ± 0.09 | 78.6 ± 2.2 |
| **SBP-7501\*** |  |  | 4 ± 0 | 118 ± 34 | 3.97 ± 0.34 | 4.79 ± 0.18 | 52.7 ± 1.1 |
| **SBP-5147\*** |  |  | 2 ± 0 | 53 ± 41 | 12.76 ± 0.14 | 14.43 ± 0.12 | 61.3 ± 0.3 |
| **6a** | Cl |  | 153 ± 14 | > 3000 | 3.65 ± 0.07 | 2.31 ± 0.09 | 95.8 ± 1.3 |
| **6b** | Br |  | 141 ± 50 | > 3000 | 5.60 ± 0.23 | 4.13 ± 0.02 | 84.8 ± 0.3 |
| **6c** | CF<sub>3</sub> |  | 62 ± 14 | 1338 ± 198 | 6.55 ± 0.12 | 6.50 ± 0.09 | 84.1 ± 1.1 |
| **6d** | CF<sub>3</sub> |  | 8 ± 2 | 172 ± 35 | 7.70 ± 0.43 | 3.73 ± 0.81 | 50.1 ± 0.5 |
| **6e** | Cl |  | 22 ± 7 | 1440 ± 523 | 0.15 ± 0.06 | 0.43 ± 0.34 | 75.6 ± 3.0 |
| **6f** | Cl |  | 21 ± 2 | > 3000 | 0.20 ± 0.13 | 0.48 ± 0.06 | 61.7 ± 2.3 |
| **6g** | Cl |  | 94 ± 43 | > 3000 | 6.78 ± 0.19 | 5.86 ± 0.12 | 84.8 ± 1.6 |
| **6h** | Br |  | 6 ± 1 | > 3000 | 8.08 ± 0.53 | 7.49 ± 0.46 | 67.9 ± 1.9 |
| **6i** | Cl |  | 2703 ± 1775 | > 3000 | 3.37 ± 0.38 | 2.45 ± 0.20 | 99.1 ± 3.1 |
| **6j** | Cl |  | 218 ± 55 | > 3000 | 6.05 ± 0.13 | 4.43 ± 0.33 | 59.8 ± 1.4 |
| **6k** | Cl |  | 292 ± 42 | 893 ± 454 | 5.75 ± 0.12 | 3.38 ± 0.14 | 105.3 ± 3.4 |
| **6l** | CF<sub>3</sub> |  | 38 ± 6 | 1957 ± 101 | 8.90 ± 0.17 | 7.08 ± 0.25 | 77.4 ± 2.0 |
| **6m** | Br |  | 32 ± 12 | > 3000 | 6.78 ± 0.13 | 4.28 ± 0.19 | 82.7 ± 0.9 |
| **6n** | CF<sub>3</sub> |  | 50 ± 25 | > 3000 | 6.53 ± 0.87 | 1.38 ± 0.33 | 80.4 ± 1.2 |
| **6o**<br>(SBP-1750) | Cl |  | 8 ± 1 | 50 ± 12 | 14.25 ± 0.11 | 12.01 ± 0.05 | 46.5 ± 1.3 |
| **6p** | CF<sub>3</sub> |  | 10 ± 0 | 7 ± 2 | 16.50 ± 0.15 | 15.81 ± 0.06 | 66.3 ± 2.2 |
| **6q** | H |  | 603 ± 100 | > 3000 | 5.18 ± 0.17 | 2.71 ± 0.09 | 104.0 ± 3.4 |
| **6r** | CF<sub>3</sub> |  | 15 ± 1 | 19 ± 3 | 17.28 ± 0.06 | 16.70 ± 0.05 | 73.3 ± 2.0 |
| **6s** | Cl |  | 38 ± 4 | 180 ± 9 | 13.28 ± 0.40 | 11.50 ± 0.27 | 78.4 ± 1.9 |
| **6t** | Br |  | 8 ± 1 | 59 ± 6 | 16.83 ± 0.12 | 16.23 ± 0.15 | 80.6 ± 2.8 |

\* Reference data for **SBI-0206965**, **SBP-7455**, and **SBP-7501**, and **SBP-5147** from Cosford et al., 2025 <sup>32</sup>

New analogues were synthesized in two steps, as outlined in **Supplemental Scheme 1**, via two consecutive substitution reactions on the pyrimidine scaffold using either **Method A** or **B**. **Method A** was employed for all analogues except those bearing a 5-trifluoromethyl substituent. Briefly, commercially available pyrimidine derivatives (**1a** or **1b**) were reacted with 2-amino-N-methylbenzamide in the presence of potassium carbonate to afford intermediates **3a** or **3b**. These intermediates were then reacted with selected amines to yield the desired ULK inhibitors. In **Method B**, 2,4-dichloro-5-(trifluoromethyl)pyrimidine (**4**) was reacted with selected amines in the presence of zinc chloride to form intermediate **5**, which underwent a second substitution with 2-amino-N-methylbenzamide at elevated temperatures to furnish the final target compounds.

Newly synthesized compounds were evaluated for ULK1 and ULK2 biochemical inhibition using a previously established ADP-Glo kinase assay ^30^. In addition to the ULK biochemical assays, compounds were also profiled in protein thermal shift (PTS) assays against the kinase domains of ULK1 and ULK2 ^30^. The data are summarized in **Table 1**.

To initiate our studies, we synthesized NVPTAE-22 (**6b**) as a benchmark compound. In addition to **6b**, we prepared the 5-chloro analogue (**6a**) and the 5-trifluoromethyl analogue (**6c**). All three compounds were evaluated for ULK1 and ULK2 inhibitory activity using the ADP-Glo assay. Compound **6a** exhibited IC₅₀ values of 153 nM and > 3000 nM against ULK1 and ULK2, respectively. Introduction of a 5-bromo group (**6b**) resulted in similar ULK1 potency (141 nM) relative to **6a**, while the 5-trifluoromethyl analogue **6c** showed nearly a 2-fold improvement in ULK1 potency (62 nM). However, all three compounds exhibited a marked reduction in ULK1 inhibitory potency compared to **SBP-7501**, which demonstrated superior biochemical activity with IC₅₀ values of 4 nM for ULK1 and 118 nM for ULK2 ^32^. Prompted by these findings and to further examine this structural modification, we synthesized compounds **6d** and **6e** by introducing the 6-amino-3,4-dihydroquinolin-2(1H)-one moiety from **SBP-7501** as the C-2 substituent. As expected, both analogues showed improved biochemical potency against ULK1 and ULK2 in the ADP-Glo assay compared to **6a** and **6c**. However, no significant enhancement in ULK1 potency was observed relative to **SBP-7501**. Notably, the 5-trifluoromethyl analogue **6d** exhibited a substantial improvement in ULK2 biochemical potency over the 5-chloro analogue **6e** (IC₅₀ = 172 nM vs. 1,440 nM), achieving potency comparable to that of **SBP-7501** (118 nM). When the 6-amino-3,4-dihydroquinolin-2(1H)-one moiety in compound **6e** was replaced with a 6-membered 1,4-benzodioxan-6-amine, an amine previously explored in our assays, the resulting compound **6j** was less potent than **6e** (IC_50_ values 218 nM and 22 nM respectively in the ULK 1 ADP-Glo assay). (**Table 1**). The analogue in which the 6-amino-3,4-dihydroquinolin-2(1H)-one moiety was replaced with a five-membered heteroaromatic ring (compound **6f**) exhibited comparable ULK1 biochemical potency (21 nM) to compound **6e** but showed a substantial decrease in activity in the ULK2 ADP-Glo assay. Other analogues within the five-membered heteroaryl series (**6g**–**6i**) did not enhance ULK2 biochemical activity, while their ULK1 potencies ranged from 6 nM to 2.7 µM, with compound **6h** being the most potent (5.7 nM). In conclusion, none of the analogues containing benzo-fused amines improved ULK biochemical potency compared to **SBP-7501**, however, several of these analogues demonstrated enhanced ULK1 potency relative to **SBI-0206965**. Next, were synthesized analogues **6k**–**6t** by replacing benzo-fused amines with substituted anilines. Among the 5-methoxy-2-methylaniline derivatives, the 5-chloro analogue **6k** demonstrated slightly improved potency in the ULK2 biochemical assay compared to the 5-trifluoromethyl (**6l)** and 5-bromo (**6m**) analogues. In contrast, **6k** showed significantly reduced ULK1 inhibitory activity (IC₅₀ = 292 nM) relative to **6l** (IC₅₀ = 38 nM) and **6m** (IC₅₀ = 32 nM). However, these compounds were still much less potent than **SBP-7501.** Interestingly, analogues **6o (SBP-1750)** and **6p**, incorporating a 3,4,5-trimethoxyaniline moiety as in our chemical probe **SBI-0206965** demonstrated improved potency in both ULK1 and ULK2 biochemical assays. In the ULK1 ADP-Glo assay, compounds **6o** and **6p** showed slightly reduced potency, with IC₅₀ values of 8 nM and 10 nM, respectively, compared to **SBP-7501** (IC₅₀ = 4 nM). Notably, the 5-chloro-substituted analogue **6o (SBP-1750)** exhibited a modest (<2-fold) improvement in ULK2 potency (IC₅₀ = 50 nM), whereas the 5-CF₃-substituted analogue **6p**, demonstrated a significant (<15-fold) improvement in the ULK2 assay (IC₅₀ = 7 nM) relative to **SBP-7501** (IC₅₀ = 118 nM). Among the 3,4,5-trimethoxyaniline derivatives **(6o–6q)**, **6q** exhibited significantly reduced potency compared to **6o** and **6p** in both the ULK1 and ULK2 assays (IC₅₀ = 603 nM, IC₅₀ = 30000 nM). Introduction of 3,5-di(morpholin-4-yl)aniline (**6r-6t**)) also resulted in similarly improved potency for ULK2 and slightly reduced ULK1 potency compared to **SBP-7501.**

Additionally, compounds were tested in the protein thermal shift (PTS) assay to verify their in vitro binding capacity and assess their relative ability to stabilize ULK 1 and ULK2, as measured by a shift in protein melting temperatures (*T_m_*) (**Table 1**). In this assay, generally, the most potent compounds in the ULK ADP-Glo assay **6o (SBP-1750)**, **6p**, **6r**, and **6t** show a larger ΔTm in the PTS assay (**Table 1**). Next, selected compounds were subsequently evaluated for their cellular target engagement with ULK1 and ULK2 using the NanoBRET assay. The results are summarized in **Table 2**. Among the compounds evaluated, four analogues; **6o** (**SBP-1750**), **6p, 6r** and **6l** exhibited improved cellular activity in the ULK1 NanoBRET assay relative to earlier analogues. **6o** (**SBP-1750**) was the most potent, with an IC₅₀ value of 4.1 nM. As shown in **Table 2**, except **SBP-7501**, all tested compounds showed relatively less ULK2 cellular potency in the NanoBRET assay.

**Table 2.** IC_50_ Values Measured by NanoBRET Assays for Selected Compounds against ULK.

| Compound | ULK1 NanoBRET IC<sub>50</sub> (nM) |
| --- | --- |
| **SBI-0206965** | 785 ± 26 |
| **SBP-7455** | 328 ± 43 |
| **SBP-7501** | 167 ± 31 |
| **SBP-5147** | 47 ± 11 |
| **6d** | 103 |
| **6l** | 3093 |
| **6o** | 4.05 |
| **6p** | 10 |
| **6r** | 17 |
| **6l** | 1.7 |

Analysis of the data presented in **Tables 1 and 2** suggests that replacing the 2-hydroxy N-methylbenzamide moiety in **SBI-0206965** with the corresponding 2-amino N-methylbenzamide group results in analogues with improved cellular and biochemical potency in both ULK1 and ULK2 assays and a large ΔTm in the PTS assay. This observation aligns with our previous findings, where substitution of an ether group with an amino group at the C-4 position of the pyrimidine scaffold enhanced ULK potency and target engagement. Based on the data from **Tables 1 and 2**, compound **6o** (**SBP-1750**) was selected for further characterization studies.

### Validation of ATG13 Knockdown as a Critical Driver of Cancer Cell Death in vitro

Previous studies from our lab demonstrate loss of ATG13 protein following ULK inhibitor treatment ^32^. We directly examined the impact of depletion of ULK1 or ATG13 in a panel of cancer cell lines, including KRAS mutant pancreatic and lung cancer cells. We observed a significant reduction in cancer cell growth and viability for both ULK1 and ATG13, with ATG13 shRNA showing stronger effects than ULK1 shRNA, perhaps reflective of redundancy between ULK1 and ULK2 in some cell lines (**Figure1-B**). These results suggest that ATG13 serves as a critical protein within the autophagy pathway and is essential for the survival of cancer cells. Given that ULK1 activation requires its interaction with ATG13, these findings indicate that developing ULK inhibitors which degrade ATG13 may be a more effective therapeutic strategy than inhibiting ULK kinase activity alone.

To further validate the role of ULK mediated autophagy in pancreatic cancer, we performed CRISPR-Cas9 knockout of key autophagy genes in MiaPaCa-2 and PANC-1 cells. Cells were cultured in soft agar for three weeks to assess anchorage-independent growth. All knockouts led to a significant reduction in colony formation compared to wild-type controls, indicating that core autophagy components are required for PDAC cell growth in this context. Notably, dual knockout of ULK1 and ULK2 resulted in the most profound growth inhibition, supporting our hypothesis that ULK are critical for tumorigenic growth (**Figure 1C**).

**Figure 1:**
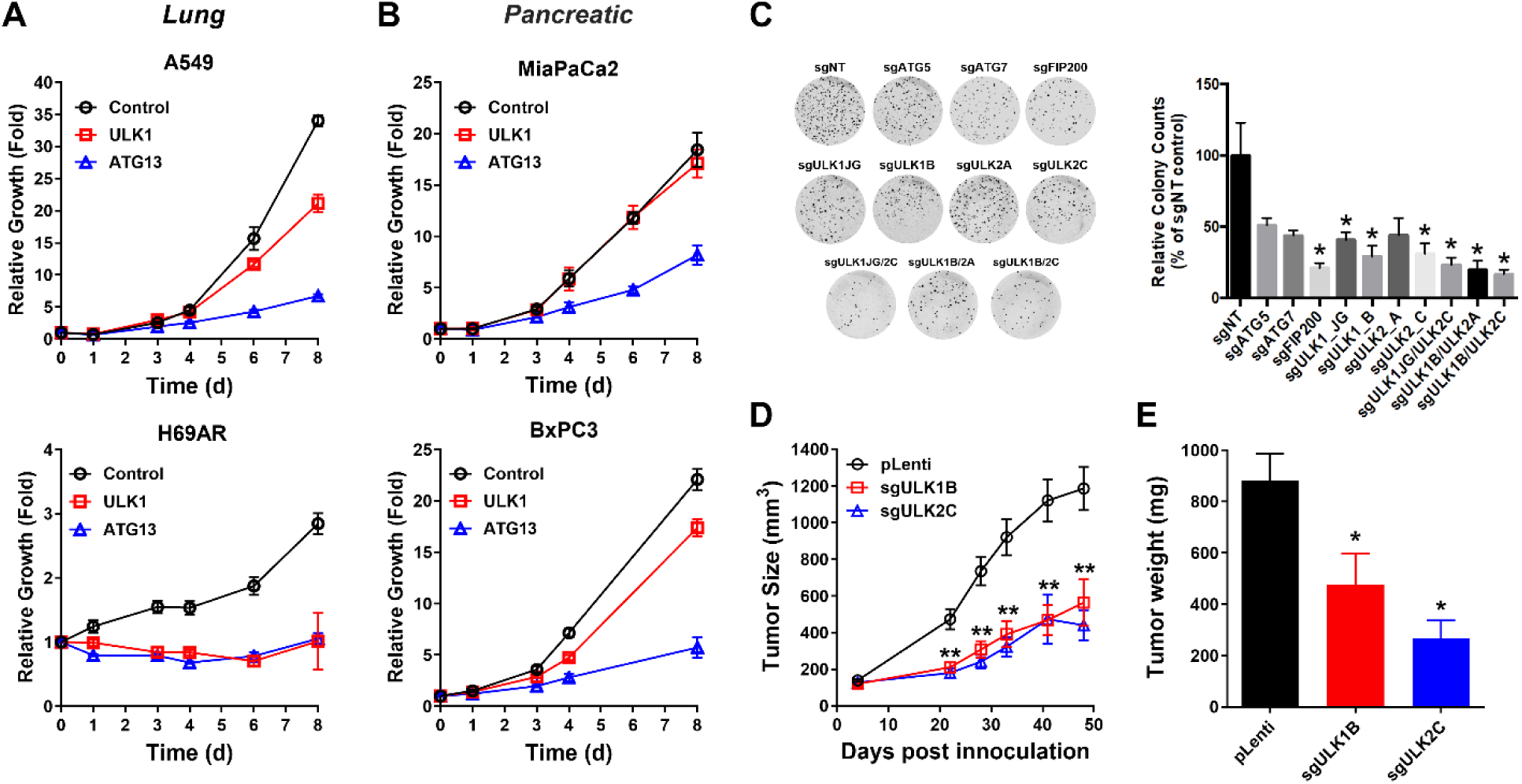
Loss of ULK1 or ATG13 lead to reduced survival of pancreatic and lung tumors. **A,B)** shRNA-mediated knockdown of ATG13 or ULK1 reduces cell viability in autophagy-dependent human cancer cell lines: A) lung and B) pancreatic. Cell viability was measured 8 days post-selection and normalized to scrambled shRNA control **C)** CRISPR-mediated knockout of essential autophagy genes reduces anchorage-independent growth in soft agar in MiaPaCa-2 pancreatic cancer cells. Cells were cultured for 3 weeks post-transduction, and colony growth was quantified relative to non-targeting control cells. **D)** Tumor growth over time following subcutaneous xenograft of CRISPR-edited MiaPaCa-2 cells into immunodeficient mice. Significance relative to sgNT control was determined by one-way Anova (p>0.0002) and t-test (*=p<0.05). **E)** Tumor volume at endpoint for mice in D). Outliers were identified using ROUT outlier analysis with Q value of 1%, Significance determined relative to sgNT control tumor size by two-way Anova (p<0.001, time x genotype interaction) followed by one-way Anova (p<0.0001) and t-test (**p<0.01) per timepoint.

To assess the contribution of ULK kinases to tumor growth in vivo, MiaPaCa-2 ULK1-or ULK2-knockout (KO) lines were implanted subcutaneously into the flanks of nude mice and tumor progression was monitored longitudinally (**Figure 1D**). Genetic loss of either ULK1 or ULK2 reduced xenograft growth and endpoint tumor mass relative to controls (**Figure 1D–E**). Immunoblotting of tumor lysates confirmed ULK1 ablation and revealed decreased phosphorylation/characteristic mobility shifts of direct ULK substrates in sgULK1 tumors, together with increased LC3B-II—the lipidated form of the autophagosome marker LC3B (MAP1LC3B), which rises when autophagosome turnover is impaired—consistent with reduced autophagic flux (**Supplemental Figure 2**). Although ULK2-KO tumors retained ULK1, they displayed reduced ATG13 phosphorylation and modulation of LC3B as well as NCOA4, a selective cargo receptor for ferritin that mediates ferritinophagy and reports on ULK-dependent selective autophagy (Supplemental **Figure 2**). Collectively, these data indicate that loss of ULK1 or ULK2 compromises autophagy and constrains pancreatic tumor growth in vivo.

### Creation of CRISPR Tagged ATG13 HiBiT A549 Cells

Our results support a critical role for ULK and ATG13 in pancreatic cancer growth and suggest that ULK inhibitors which degrade ATG13 may have enhanced therapeutic potential. To identify ATG13 degraders we established an assay to reliably measure cellular levels of ATG13 protein in a high throughput manner. ATG13 was endogenously tagged with the small 11-amino-acid (1.3 kDa) luminescent HiBiT peptide (Promega) at the end of the final exon, exon 18, but before the stop codon of ATG13 ^33^. The small size of the tag and location of insertion allow for the study of ATG13 in a nearly native state. To create this endogenous ATG13 HiBiT tag, we used ribonucleoprotein (RNP) complexes consisting of CRISPR/Cas9 and tracrRNA/crRNA targeting the 3’ end of ATG13. A549 cells were electroporated with this RNP in the presence of a dsDNA homology-directed repair template containing the HiBiT sequence, flanked by homology arms matching the area surrounding the cut site. A549 cells are an ATG-dependent, KRAS-driven NSCLC cell line, making them a suitable model for autophagy inhibitor studies ^34–36^. When ATG13 is successfully tagged, the HiBiT peptide complements with the LgBit subunit of NanoLuc added to the cells post-treatment, generating a luminescent signal measurable by a standard bioluminescent plate reader (**Figure 2A**).

**Figure 2:**
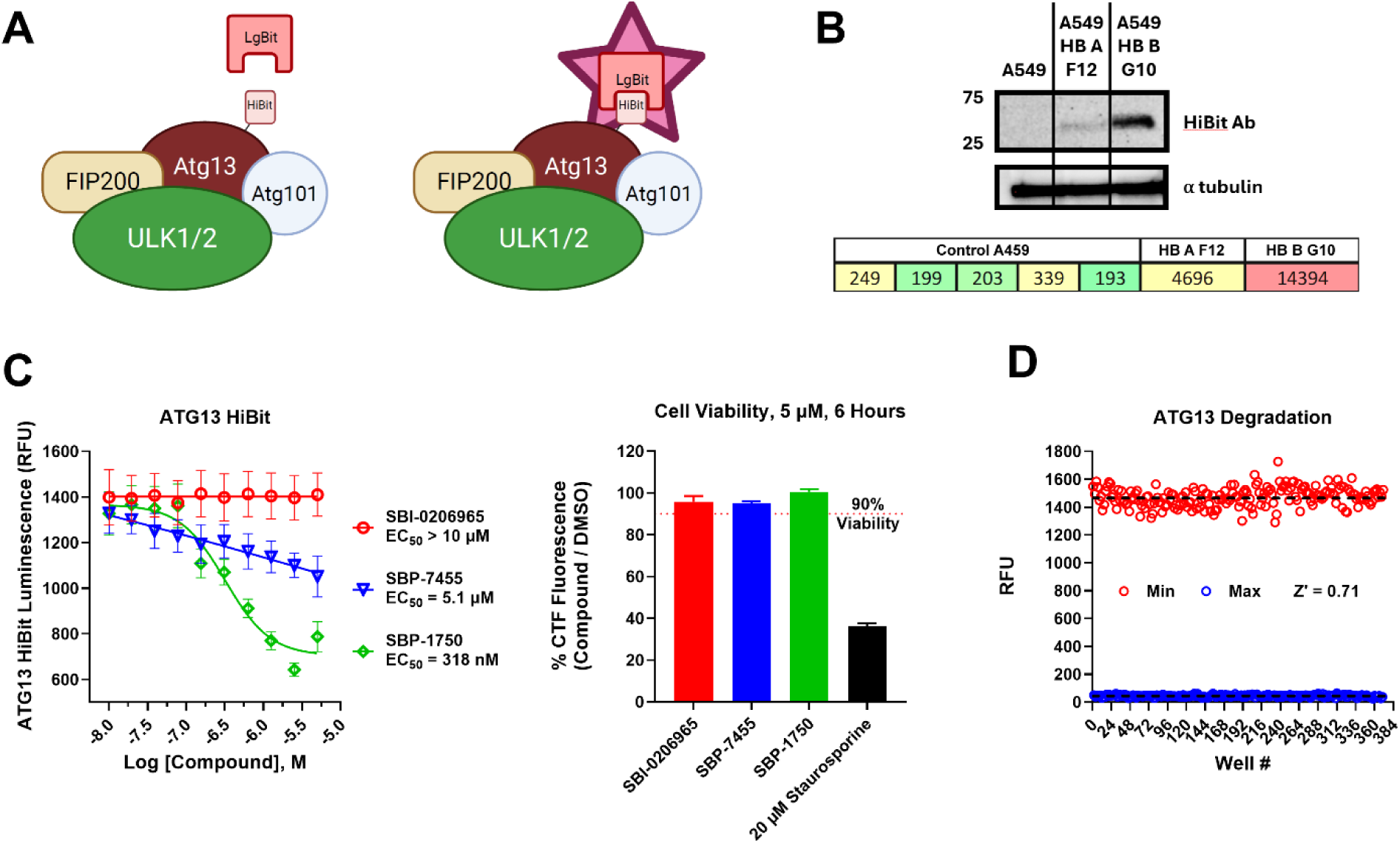
Development of an ATG13 HiBiT cell-based in vitro assay in A549 cells. **A)** Diagram of a HiBiT tag on ATG13 interacting with LgBit reagent, creating a bright luminescent signal. **B)** Confirmation of insertion of HiBiT tag onto ATG13 in A549 Cells. Western Blot with HiBiT antibody shows band at 72 kDa (expected size of ATG13). HiBiT Lytic Assay performed with selected clones and A549 cells without CRISPR editing. **C)** Optimizing the ATG13 HiBiT Lytic Assay. 5 µM compound treatment for 6 hours showed ATG13 degradation but minimal cell death. **D)** Calculating the z’ factor of 384 well ATG13 HiBiT Lytic Assay. Z’ = 0.71 Following electroporation and recovery, we performed single-cell sorting to isolate a monoclonal population of ATG13 HiBiT-tagged A549 cells. These clones were validated using the Promega HiBiT lytic detection system, which showed that two wells exhibited a strong bioluminescent signal, especially in comparison to control A549 cells (**Figure 2C**). Additionally, a HiBiT-specific antibody was used in a Western blot to confirm successful tagging of ATG13 (72 kDa) (**Figure 2B**).

Once the HiBiT tag was successfully incorporated into ATG13 in A549 cells, we developed an assay to measure ATG13 degradation in response to ULK inhibitors. We optimized time points (4, 6, 16, 24, and 48 hours) and doses to identify conditions where ATG13 degradation could be observed without significant toxicity. We observed significant ATG13 degradation at 5 µM with a 6-hour treatment, while maintaining less than 10% loss in cell viability, enabling us to distinguish ATG13 degradation from toxicity (**Figure 2C**). We have further optimized the ATG13 HiBiT assay to be suitable for high-throughput screening in a 384-well format, achieving a robust Z’ factor of 0.71 (**Figure 2D**).

### SBP-1750 Is the Most Potent ATG13-Degrading ULK Inhibitor

To systematically evaluate the impact of ULK inhibitors on ATG13 degradation, we. screened a panel of structurally related ULK inhibitors, including those developed within our lab as part of the SAR/chemistry studies detailed earlier in this paper, as well as externally developed ULK inhibitors. Among all compounds tested, **SBP-1750** emerged as the most potent inducer of ATG13 degradation, exhibiting significantly stronger activity than any other ULK inhibitor.

Notably, within our internal compound series, **SBP-1750**, our current clinical lead, exhibited the strongest ATG13 degradation, surpassing **SBP-7501** and **SBP-5147**, our recently published in vivo tool compounds, and **SBP-7455**, an earlier generation chemical probe ^32^. Potency in ATG13 degradation assays strongly correlated with ULK inhibitory activity and binding affinity. Consistently, externally developed ULK inhibitors tested in this study failed to induce substantial ATG13 degradation, further supporting that this effect is a unique property of the structural scaffolds designed in our lab (**Figure 3A**).

**Figure 3:**
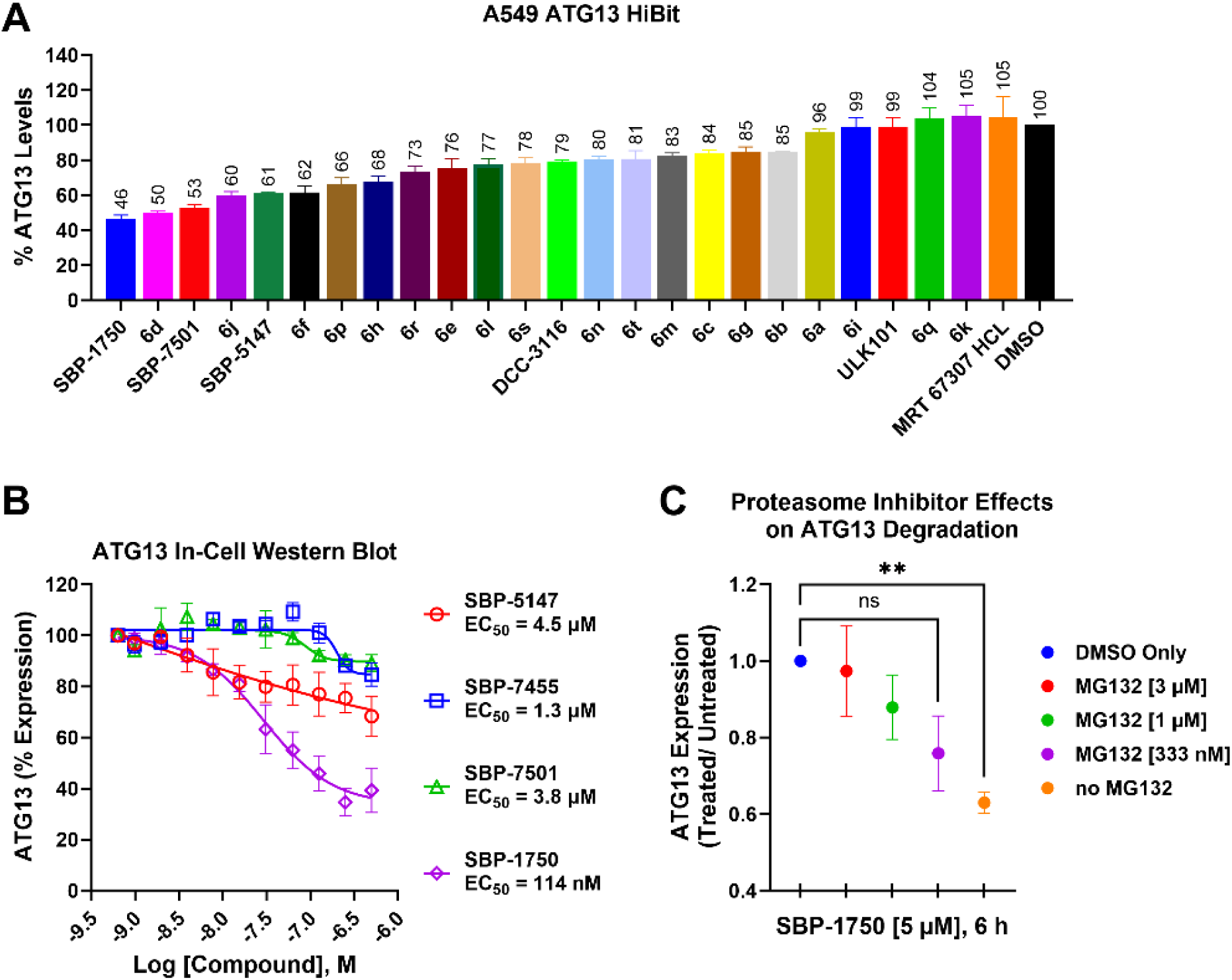
SBP-1750 induces the largest ATG13 degradation in A549 cells. **A)** HiBiT-based lytic assay quantifying ATG13 degradation in A549 cells following 6-hour treatment with 5 µM of novel ULK inhibitors and representative ULK inhibitors from the literature. Luminescence reflects loss of HiBiT-tagged endogenous ATG13. **B)** Validation of ATG13 degradation by In-cell Western blot in A549 ATG13-HiBiT cells treated for 48 hours with selected compounds **C)** ATG13 degradation in A549 ATG13-HiBiT cells is rescued by proteasome inhibition. Cells were treated for 6 hours with DMSO (blue), compound only (orange, 5 µM), or compound (5 µM) in combination with varying concentrations of MG132. HiBiT luminescence was used to quantify ATG13 levels. Significance calculated via one-way Anova with a post-hoc Dunnett test. ** p<0.005.

### Confirming ATG13 Degradation via In Cell Western Blot

To validate the results of the ATG13 HiBiT degradation screen with our ULK inhibitors, we employed LI-COR In-Cell Western blots (ICWBs) as a secondary assay. This quantitative assay allows for the visualization and measurement of protein expression directly in fixed cells using infrared fluorescence-conjugated secondary antibodies. ICWBs provide a robust and reproducible method for quantifying ATG13 levels in a dose-dependent manner while allowing for normalization to control proteins to account for any variation in cell number. Treatment with SBP-1750 induced ATG13 degradation with an IC50 of 114 nM as measured by ICWB (**Figure 3B**). These data are consistent with our previous results and confirm the predictive value of the ATG13 HiBiT screen.

### Mechanistic Evaluation of ATG13 Degradation

The two primary mechanisms of intracellular protein degradation are autophagy and the ubiquitin proteasome system^37^. To investigate the mechanism responsible for ATG13 degradation induced by our ULK inhibitors, we co-treated cells with 5 µM of the top ATG13 degrading compound, **SBP-1750**, for 6 hours (mirroring conditions in the ATG13 HiBiT screen) alongside the proteasome inhibitor MG132 at varying doses. Our results demonstrated that co-treatment with MG132 significantly altered the level of ATG13 degradation compared to treatment with the ULK inhibitor alone (**Figure 3C**). This marked reduction in ATG13 degradation upon proteasome inhibition suggests that the degradation may be proteasome-dependent, indicating that our compound may induce ATG13 degradation through a proteasomal pathway, suggesting that our compound is working similarly to a molecular glue on ATG13 ^38^.

### ATG13-Degrading Compounds Are the Most Potent Autophagy Inhibitors

We aimed to evaluate whether the ATG13 degradation observed with our lead ULK inhibitors also reflected inhibition levels, utilizing A549 cells engineered to express the GFP-LC3-RFP-LC3ΔG construct ^20^. This assay enables dynamic monitoring of autophagic flux by tagging LC3 with both GFP and RFP, fluorescent markers that behave differently in autophagic and lysosomal environments. GFP fluorescence is sensitive to the acidic environment within autolysosomes and is quenched upon LC3 degradation, while RFP remains stable, allowing reliable differentiation between autophagosomes and autolysosomes ^20^. Cells were incubated in Earle’s balanced salt solution (EBSS) and treated for 18 hours with each compound at a concentration of 5 µM. The use of EBSS in autophagic flux assays provides a robust and standardized induction of ATG, creating the dynamic range necessary to evaluate the effects of inhibitors in cancer cells ^39^. Our findings indicate that ULK inhibitors which also degrade ATG13 robustly inhibit autophagy (**Figure 4A**). Specifically, the compound **SBP-1750**, which exhibited the greatest ATG13 degradation, showed the highest level of autophagic inhibition in the GFP-LC3-RFP-LC3ΔG assay (**Figure 4A**). This correlation suggests that ATG13 degradation, as measured by our HiBiT assay, may serve as a reliable indicator of a compound’s ATG-inhibitory potential. The consistency across these two assays supports the notion that ATG13 degradation levels could reflect downstream effects on ATG, potentially offering a useful metric for predicting autophagy inhibition among ULK inhibitors.

**Figure 4:**
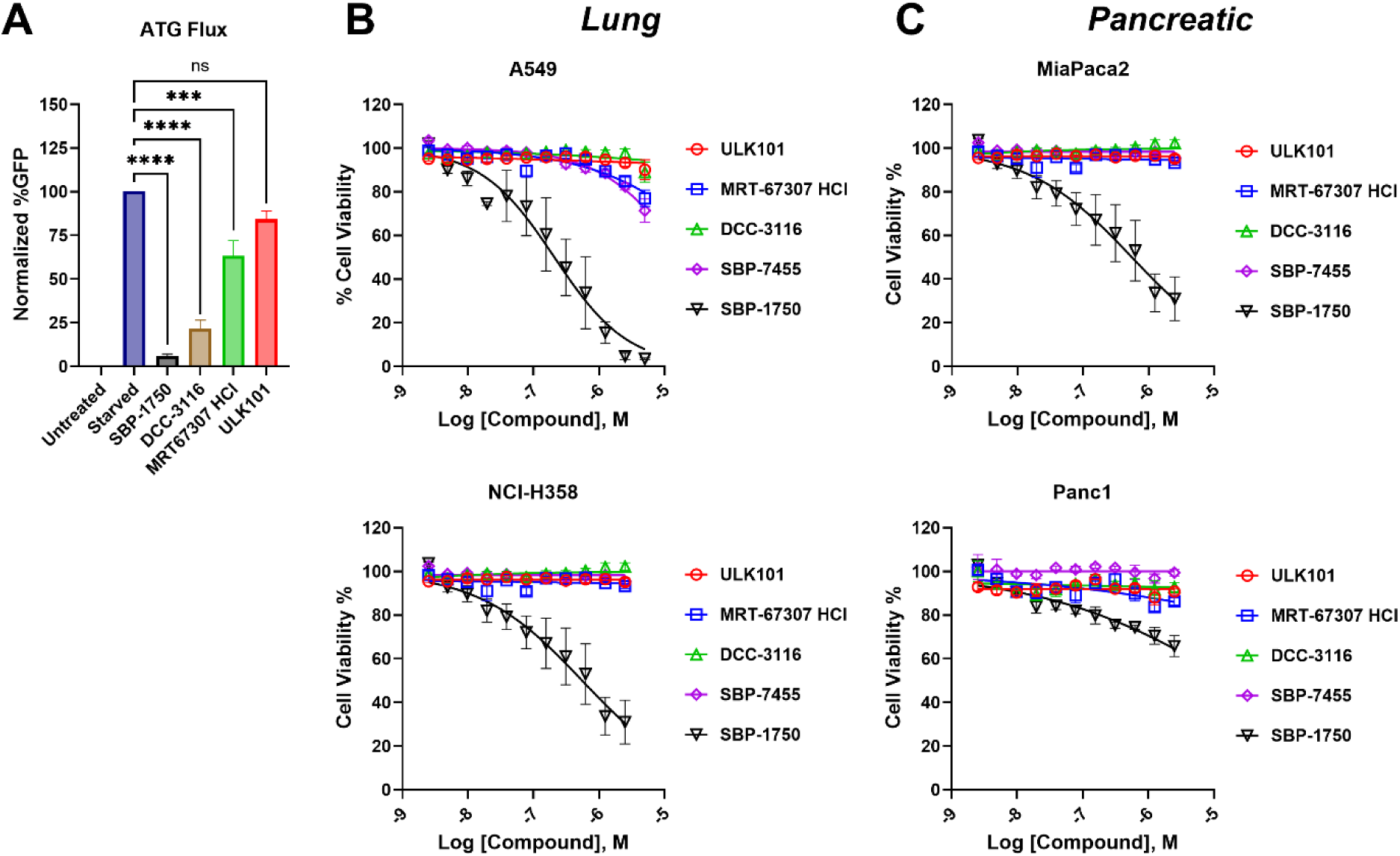
ATG13 Degradation Correlates with Autophagy Inhibition and Cancer Cell Death. **A)** Autophagic flux measured using the GFP-LC3–RFP-LC3ΔG reporter in A549 cells treated with 5 µM ULK1/2 inhibitors for 18 hours under starvation conditions (EBSS). Significance was determined using a one-way ANOVA with a post-hoc Tukey test. *** p<0.001, **** p<0.0001. Autophagy Flux was normalized to untreated controls (100% autophagy Flux) to determine relative inhibition of autophagy. Significance calculated via One Way Anova. **B and C)** Cell viability measured by CellTiter-Glo in a panel of KRAS G12-mutant cancer cell lines treated with increasing concentrations of ULK1/2 inhibitors for 48 hours. Viability was normalized to vehicle-treated controls.

### ATG13-Degrading and ATG-Inhibiting Compounds Induce the Greatest Cell Death in KRAS Mutant Cancer Cells

Given the substantial evidence that autophagy supports the survival of KRAS driven cancers ^40, 41^, we sought to determine whether compounds that induced the most significant ATG13 degradation, and subsequently the highest levels of autophagy inhibition, would also correlate with increased tumor cell death. To test this, we employed a CellTiter-Glo cell viability assay, treating KRAS mutant cancer cells with varying doses of each compound. Consistent with our hypothesis, **SBP-1750** showed the highest reduction in cell viability, mirroring the ranking observed in both

ATG13 degradation and autophagy inhibition (**Figure 4B–C**). These findings underscore the potential therapeutic significance of ATG13 degradation and autophagy inhibition in driving cell death in tumor cells and highlight **SBP-1750** as a particularly potent inhibitor in this context.

### In Vivo PK of SBP-1750

To evaluate the PK properties and oral availability of **SBP-1750**, we administered a single dose (10 mg/kg) via oral gavage to mice and assessed plasma exposure over 24 h by LC-MS/MS. The PK analysis of **SBP-1750** (**Figure 5**) showed that the time to peak plasma concentration (T_max_) was 1 hour and the maximum plasma concentration (C_max_) was 32.9 µM, a value approximately more than 8,000-fold higher than its NanoBRET IC_50_ value for ULK1 (IC_50_= 4.05 nM) and more than 100-fold higher than its NanoBRET IC_50_ value for ULK2 (IC_50_= 230 nM). Moreover, the plasma concentration of SBP-1750 remained above the ULK NanoBRET IC_50_ value for at least 8 h post-dosing with a T_1/2_ of 2 h (**Figure 5**). These data indicate that the plasma levels of **SBP-1750** are sufficient for in vivo target engagement. Compared to **SBP-5147**, which had a C_max_ of 664 nM and T_max_ of 0.25 h when dosed at 10 mg/kg orally ^30, 32^, **SBP-1750** presented improved PK properties with higher C_max_ and AUC levels, reflecting increased in vivo exposure. Based on this favorable PK data, **SBP-1750** was selected for in vivo efficacy studies.

**Figure 5:**
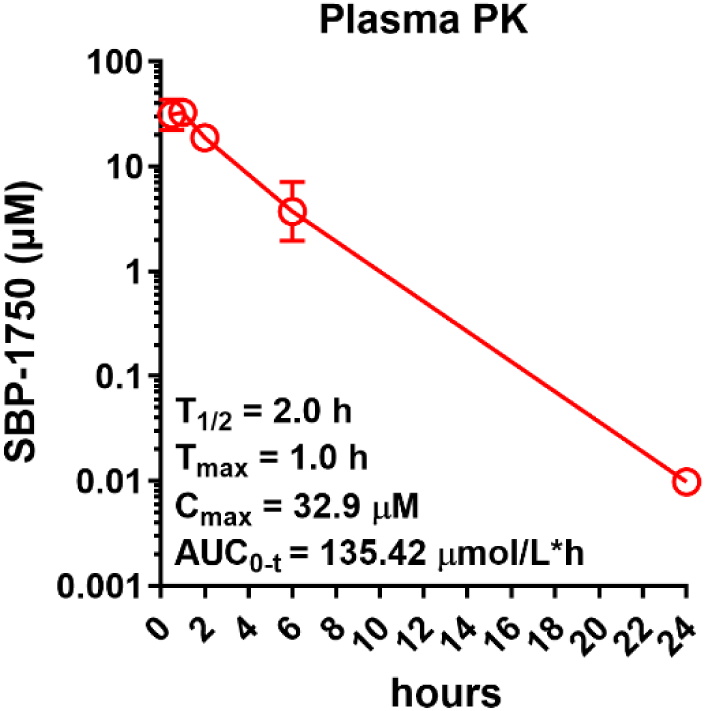
PK Profile of SBP-1750 Following Oral Dosing in C57BL/6 Mice. 24 h time-course concentrations of **SBP-1750** measured following 10 mg/kg oral dosing (1 mg/mL, 10 mL/kg). Compound was formulated in 5% DMSO, 10% Tween-80, and 85% water. Data shown are N=3 and represent the geometric mean ± geometric SD.

### SBP-1750 Exhibits Strong Tolerability, Reduces Pancreatic Tumor Size, and Reduces Metastasis in Mice

To investigate the in vivo efficacy of compound **SBP-1750** in pancreatic cancer, we established orthotopic syngeneic tumors using the murine pancreatic ductal adenocarcinoma (PDAC) cell line KPC4662 in immunocompetent C57BL/6 mice. The KPC4662 cell line is a murine PDAC model derived from the well-established KPC mouse model, which harbors pancreas-specific mutations in *Kras*^G12D^ and *Trp53*^R172H^ ^42^. Mice were dosed with either **SBP-1750** at 40 mg/kg or vehicle beginning on day 10 post-implantation by oral gavage, with peripheral blood collection performed on days 12 and 27. Tumors were harvested on day 28 for endpoint analysis (**Figure 6A**). PD analysis of tumor tissue revealed that treatment with compound **SBP-1750** led to a marked reduction in ATG13 protein levels (**Figure 6B**), consistent with the ULK complex degradation observed in vitro (**Figure 3C**). This confirms that SBP-1750 effectively penetrates the tumor and engages its intended target in vivo, disrupting autophagy initiation in part through ATG13 degradation and ULK complex destabilization. Throughout the 17-day treatment period, mice treated with compound **SBP-1750** exhibited no significant differences in body weight compared to vehicle-treated controls, indicating acceptable tolerability (**Figure 6C**).

**Figure 6:**
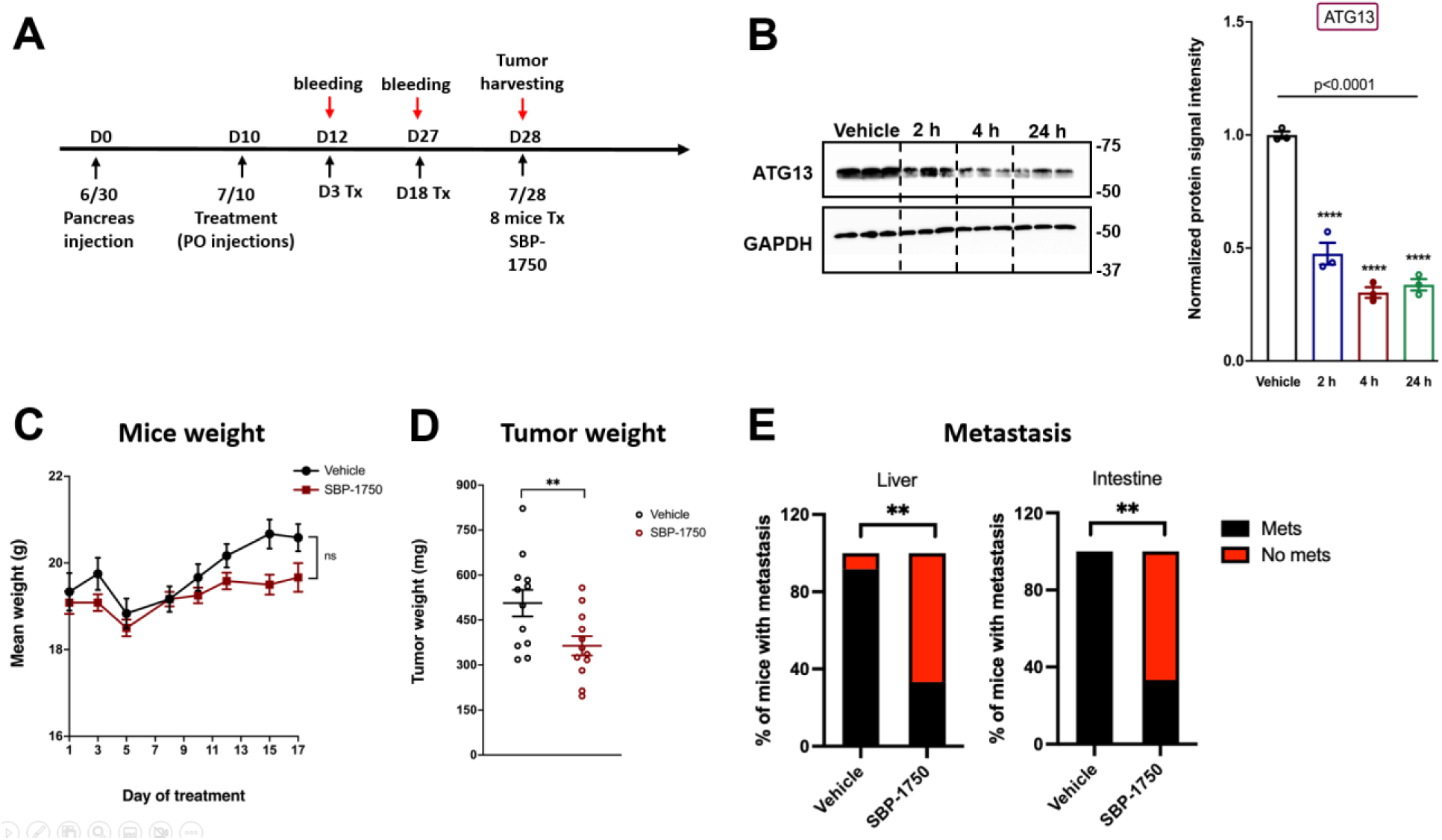
SBP-1750 Reduces Pancreatic Tumor Size and Reduces Metastasis in the KPC4662 PDAC Mouse Model. **A)** Treatment and harvesting schedule for in vivo experiments. n = 12 vehicle treatment, n = 12 **SBP-1750** treatment. **B)** Representative Western blot (left) and PD analysis (right) of ATG13 levels in tumor tissue collected from treated mice. **C)** Weight of mice over the course of treatment with either vehicle (DMSO) or **SBP-1750 D)** Tumor weights of KPC orthotopic tumors treated with vehicle or 40 mg/kg **SBP-1750** (n = 12)**. E)** Tumor macrometastases to the liver and intestine were analyzed. The percentage of mice with evidence of macrometastases to the liver and intestine was determined. Chi-squared tests. **p<0.01.

Additionally, endpoint tumor weights were significantly reduced in the **SBP-1750**-treated group, demonstrating potent antitumor efficacy (**Figure 6D**). Furthermore, mice treated with **SBP-1750** displayed a significant reduction in metastatic burden to both the liver and intestines relative to vehicle-treated animals, suggesting that **SBP-1750** may inhibit both primary tumor growth and metastatic dissemination in this model (**Figure 6E**).

### SBP-1750 Increases both CD4+ and CD8+ T Cell population

It is widely appreciated that autophagy can suppress immune signaling, help maintain the immunosuppressive tumor microenvironment, clear damage associated molecular patterns (DAMPs), and reduce antigen presentation ^43–45^. Previous studies demonstrate that autophagy inhibition can increase antigen presentation and improve immune checkpoint therapy ^46^. Based on these data, we evaluated the tumor microenvironment following treatment of **SBP-1750**. As part of our endpoint tumor analysis, we performed immunofluorescence staining for CD4⁺ and CD8⁺ T cells to assess immune infiltration. Tumors from mice treated with compound **SBP-1750** showed an increase in both CD4⁺ and CD8⁺ T cell populations compared to vehicle-treated controls (**Figure 7A-B**). These cell types play central roles in anti-tumor immunity: CD8⁺ cytotoxic T cells directly kill tumor cells, while CD4⁺ helper T cells coordinate immune responses and support effector cell function ^47^. The enhanced infiltration of these lymphocyte subsets suggests that **SBP-1750** may not only suppress tumor growth directly but also promote anti-tumor immune activation, highlighting its potential to synergize with immune checkpoint inhibitors in future combination therapies.

**Figure 7:**
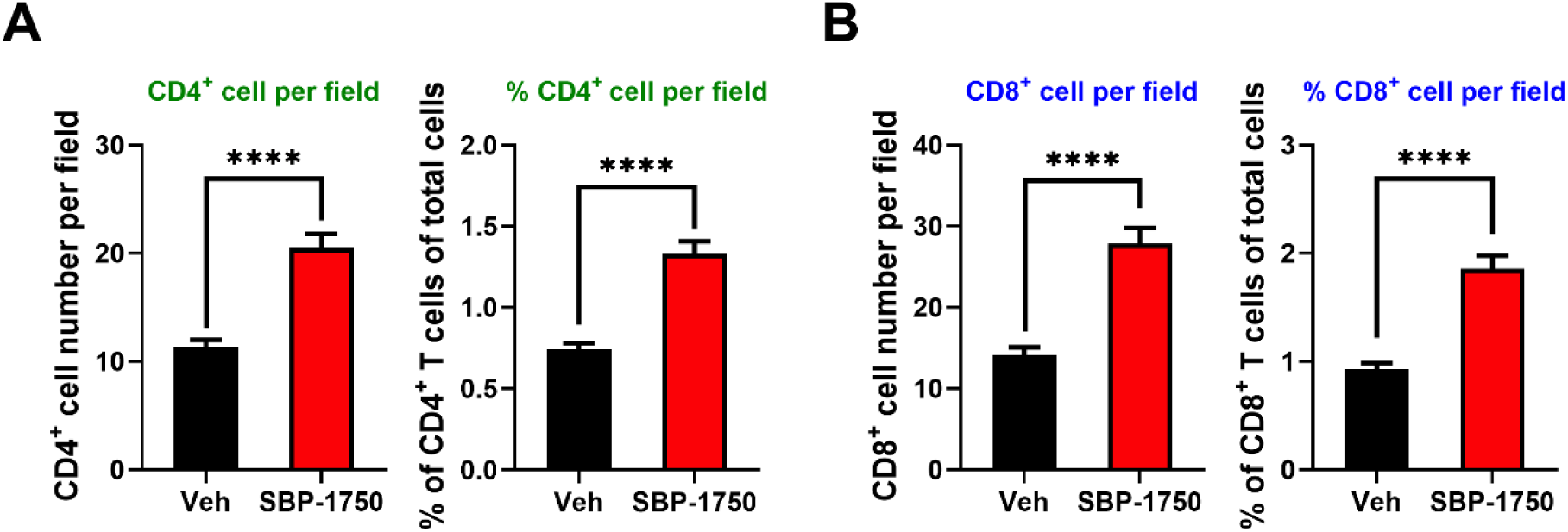
SBP-1750 increases both CD4^+^ and CD8^+^ T cell infiltration in KPC orthotopic tumors. (**A and B)** Quantification of CD4^+^ (A) and CD8^+^ (B) T cells in KPC orthotopic tumors treated with vehicle or **SBP-1750**, shown as cell number per image field or as the percentage relative to total cells in each field. Statistical significance was calculated using an unpaired two-tailed nonparametric Mann-Whitney T-test. **** p<0.0001.

## DISCUSSION

This work highlights a promising new class of ULK inhibitors and emphasizes the role of ATG13 degradation as a potential marker and therapeutic target in cancer autophagy inhibition. Autophagy is a core survival program in established cancer cells, essential in tumor growth progression and therapeutic resistance. Particularly, oncogenic KRAS elevates basal autophagy in order to maintain healthy mitochondria, sustain TCA-cycle metabolites, and support tumorigenesis ^48^. Unfortunately, despite the clinical relevance of autophagy in cancer, there are currently no approved autophagy inhibitors available to patients. Agents such as hydroxychloroquine (HCQ) and chloroquine (CQ) are lysosomal inhibitors with broad, non-specific activity, and their clinical utility as ATG-targeting agents remains limited. Only one ULK inhibitor, DCC-3116, has made it to clinical trials and is still in early-phase development (Phase 1/2) and no direct inhibitors of the autophagy initiation pathway have reached regulatory approval ^49^. However, our data suggest that DCC-3116 has little cytotoxicity as a monotherapy (**Figure 4B and C**) which is supported by previous reports ^50^. As a selective ULK inhibitor that promotes ATG13 degradation, and displays single agent activity, SBP-1750 offers distinct advantages.

While optimizing the SAR to improve drug-like properties of our ULK inhibitors, we observed a striking trend: increased ATG13 degradation coincided with improved compound profiles. Building on this observation, we describe the discovery and characterization of a novel series of ULK inhibitors with the unique phenotype of promoting ATG13 degradation through the ubiquitin proteasomal system. Through systematic testing of our lead ULK inhibitors, we identified compounds that not only effectively inhibit ULK but also induce significant ATG13 degradation, an outcome not observed with other autophagy or ULK inhibitors. Our data revealed a clear correlation: compounds that caused greater ATG13 degradation also showed enhanced autophagy inhibition and led to increased cancer cell death, suggesting that ATG13 degradation could serve as a both a predictive biomarker and a potential target for autophagy inhibition and therapeutic potential in cancer. Targeted protein degradation and molecular glues are a rapidly expanding fields within drug discovery, with several compounds being explored in clinic ^23, 25–29,51^. **SBP-1750** offers a unique advantage over other autophagy inhibitors and even other ULK inhibitors, in that it not only inhibits the active sites of the essential ULK kinases but also causes the essential adaptor protein ATG13’s degradation via the ubiquitin proteasome system like a molecular glue. This dual activity enhances **SBP-1750’s** ability to inhibit autophagy and kill tumor cells over a traditional kinase inhibitor.

**SBP-1750** is well-tolerated in C57BL/6 mice, which demonstrate no significant weight loss compared to mice treated with vehicle alone, while demonstrating single-agent efficacy to reduce tumor size, tumor weight, and metastasis in the KPC4662 pancreatic tumor model. Therefore, we have created an effective and tolerable autophagy inhibitor, whereas the FDA approved compounds chloroquine and hydroxychloroquine are associated with adverse side effects. In combination with its success as a monotherapy, this compound’s ability to inhibit ATG, an established driver of chemoresistance, supports its use in combination with standard chemotherapeutics, potentially accelerating its path to the clinic. Yet beyond these well-recognized advantages, we made an especially exciting discovery: treatment with **SBP-1750** also increased intratumoral CD4⁺ and CD8⁺ T cell levels in this pancreatic cancer model. This unexpected immune activation suggests that **SBP-1750** may also enhance responses to immunotherapies such as anti–PD-L1, positioning it as a uniquely versatile agent capable of improving outcomes across both cytotoxic and immune-based treatment strategies.

In summary, we have described the design, synthesis, and characterization of a novel autophagy inhibitor that demonstrates superior therapeutic potential compared to existing ATG-targeting agents. This compound, **SBP-1750**, inhibits autophagy by binding to and inhibiting ULK kinase activity and causing the degradation of the essential adaptor protein ATG13. The degradation of ATG13 not only serves as a robust PD biomarker but may also contribute directly to the compound’s enhanced antitumor activity. **SBP-1750** strongly inhibits ULK kinase activity, blocks ATG, degrades ATG13 both in vitro and in vivo, and significantly reduces tumor burden in mouse models while remaining well tolerated. Together, these findings establish **SBP-1750** as a compelling next-generation ULK inhibitor with the potential to advance into clinical development as a novel therapeutic strategy for targeting ATG-dependent cancers.

## METHODS

### General Chemistry Information

All reactions were performed with oven-dried glassware under a nitrogen atmosphere with magnetic stirring. All solvents and chemicals were purchased from commercial sources and used without further purification unless otherwise stated. Reaction progress was monitored by LC-MS and/or thin-layer chromatography (TLC). Chromatographic purification was carried out using prepacked silica or C18 cartridges from RediSep and eluted using an ISCO companion system. Reverse-phase purifications were conducted using water and acetonitrile doped containing 0.1% formic acid modifier. Purity and characterization of compounds were established by a combination of liquid chromatography-mass spectroscopy (LC-MS) and NMR analytical techniques and was >95% for all tested compounds. ^1^H– and ^13^C-NMR spectra were obtained on a Joel 400 spectrometer at 400 MHz and 101 MHz, respectively. Chemical shifts are reported in *δ* (ppm) relative to residual solvent peaks or TMS as internal standards. Coupling constants are reported in Hz. High-resolution ESI-TOF mass spectra were acquired from the Mass Spectrometry Core at The Sanford-Burnham Medical Research Institute (Orlando, Florida, Site closed). HPLC-MS analyses were performed on a Shimadzu 2010EV LCMS using the following conditions: Kromisil C18 column (reverse phase, 4.6 mm x 50 mm); a linear gradient from 10% acetonitrile and 90% water to 95% acetonitrile and 5% water over 4.5 min; flow rate of 1 mL/min; UV photodiode array detection from 200 to 300 nm.

#### General Procedure for the synthesis of 5-Chloro (Bromo)-N^2^,N^4^-diarylpyrimidine-2,4-diamine derivatives

*General method A*: To a solution of appropriate amine (1 equiv.) in DMF, were added 2,4,5-trichloro pyrimidine/2,4-dichloro-5-bromopyriidine (1.3 equiv.) and K_2_CO_3_ (1.3 equiv.). The reaction mixture was stirred at 75 °C for 5 h. It was then cooled to room temperature and then poured into water. The resulting precipitate was collected by filtration followed by washing with 50% aqueous acetonitrile and then dried under reduced pressure to afford the desired 2,5-dichaloro-*N-*arylpyrimidin-4-amine derivative. The crude product was used for next step without further purification.

*General method B*: To a stirred solution of the product from “General Method A” (1 equiv.) in ^n^BuOH was added appropriate aniline (2 equiv.) and heated at 110 °C for 18-24 h. The reaction mixture was cooled to room temperature and excess solvent was reduced under reduced pressure. The crude residue was purified using automated prep-HPLC to yield the desired 5-halo-*N*^2^*,N*^4^-diarylpyrimidine-2,4-diamine.

*General method C*: To a stirred solution of the product from “General Method A” (1 equiv.) in EtOH, were added appropriate amine and (1 equiv.) and HCl. The reaction mixture was stirred at 60 °C for 4-6 h. It was then cooled to room temperature diluted with water and then neutralized with 1N NaOH solution and extracted with ethyl acetate (3 times). The combined organic layers were washed with water, brine and dried over anhydrous Na_2_SO_4_. Removal off the solvent under reduced pressure afforded the crude product. The crude residue was purified using automated prep-HPLC to yield the desired 5-halo-*N*^2^*,N*^4^-diarylpyrimidine-2,4-diamine.

#### General Procedure for the synthesis of N^2^,N^4^-diaryl-5-(trifluoromethyl)pyrimidine-2,4-diamine derivatives

*General method D*: To a solution of 5-trifluromethyl-2,4-dichloropyrimidine (1 equiv.) in DCE:^t^BuOH (1:1) was added ZnCl_2_ (1.2 equiv) at 0 °C. After 1 h, appropriate aniline (1 equiv.) and triethylamine (1.2 equiv) in DCE:^t^BuOH was added to the reaction mixture. After stirring for 1.5 h, the reaction mixture was concentrated to get the crude product. The crude product was triturated with MeOH, filtered and dried to yield the desired 5-trifluromethyl-4-chloro-*N-* arylpyrimidin-2-amine derivative.

*General method E*: To a stirred solution of the product from “General Method D” (1 equiv.) in ^n^BuOH, were added appropriate aniline (1 equiv.) and DIEA (1equiv.). The reaction mixture was stirred at 110 °C for 16 h. It was then cooled to room temperature excess solvent was reduced under reduced pressure. The crude residue was purified using automated prep-HPLC to yield the desired 5-trifluromethyl-*N*^2^*,N*^4^-diarylpyrimidine-2,4-diamine.

**2-(2,5-Dichloropyrimidin-4-ylamino)-*N*-methylbenzamide (3a).** The title compound was prepared according to General Method A by the reaction of 2-amino-*N*-methyl benzamide (1.5 g, 10 mmol) in DMF, were added 2,4,5-trichloro pyrimidine (2.38 g, 13 mmol) and K_2_CO_3_ (1.79 g, 13 mmol). White solid (2.6 g, 89%).**^1^H NMR** (DMSO-d_6_): *δ* 12.15 (s, 1H), 8.82 (brs, 1H), 8.48 (d, *J* = 7.3 Hz, 1H), 8.43 (s, 1H), 7.76 (d, *J* = 7.3 Hz, 1H), 7.57 (t, *J* = 8.2 Hz, 1H), 7.18 (t, *J* = 7.8 Hz, 1H), 2.76 (d, *J* = 2.3 Hz, 3H). **^13^CNMR** (DMSO-d_6_): *δ* 169.2,156.7,155.9, 138.7, 132.4, 128.7, 123.7, 121.7, 121.4, 115.5, 27.3. LC-MS (ESI) calcd. for C_12_H_10_Cl_2_N_4_O [M+H]^+^: 297.02; found: 297.00.

**2-(5-Bromo-2-chloropyrimidin-4-ylamino)-*N*-methylbenzamide (3b).** The title compound was prepared according to General Method A by the reaction of 2-amino-*N*-methyl benzamide (3.0 g, 20 mmol), 5-bromo-2,4-dichloropyrimidine (5.5 g, 24 mmol) and K_2_CO_3_ (3.3 g, 24 mmol). Yellow solid (4.5 g, 66%).**^1^H NMR** (DMSO-d_6_): *δ* 11.93 (s, 1H), 8.50 (brs, 1H), 8.48 (d, *J* = 7.3 Hz, 1H), 8.41 (s, 1H), 7.75 (d, *J* = 7.3 Hz, 1H), 7.54 (t, *J* = 8.2 Hz, 1H), 7.17 (t, *J* = 7.8 Hz, 1H), 2.76 (d, *J* = 2.3 Hz, 3H). **^13^CNMR** (DMSO-d_6_): *δ* 169.4, 158.6, 157.9, 156.2, 139.8, 136.8, 131.9, 128.5, 122.4, 121.9, 121.4, 26.8. LC-MS (ESI) calcd. for C_12_H_10_BrClN_4_O [M+H]^+^: 340.79; found: 340.85.

**6-(4-Chloro-5-(trifluoromethyl)pyrimidin-2-ylamino)-3,4-dihydroquinolin-2(1H)-one (5a).** The title compound was prepared by the reaction of 5-trifluromethyl-2,4-dichloropyrimidine (1 g, 4.6 mmol), ZnCl_2_ (5.5 mL, 5.5 mmol), 6-amino-3,4-dihydroquinolin-2(1H)-one hydrochloride (0.841 g, 4.6 mmol) and triethylamine (1 g, 5.5 mmol) according to general method D. Pale yellow solid (1 g, 64%) as a yellow solid. **^1^H NMR** (DMSO-d_6_): *δ* 10.51 (s, 1H), 10.04 (s, 1H), 8.79 (s, 1H), 7.40 (s, 1H), 7.39 (d, *J* = 8.2 Hz, 1H), 6.82 (d, *J* = 12.6 Hz, 1H), 2.76 (t, *J* = 7.2 Hz, 2H), 2.39 (t, *J* = 7.3Hz, 2H). LC-MS (ESI) calcd. for C_14_H_10_ClF_3_N_4_O [M+H]^+^: 343.04; found: 342.90.

**4-Chloro-5-(trifluoromethyl)-*N*-(3,4,5-trimethoxyphenyl)pyrimidin-2-amine (5b).** The title compound was prepared by the reaction of 5-trifluromethyl-2,4-dichloropyrimidine (1 g, 4.6 mmol), ZnCl_2_ (5.5 mL, 5.5 mmol), 3,4,5-trimethoxyaniline (0.841 g, 4.6 mmol) and triethylamine (1 g, 5.5 mmol) according to general method D. Pale yellow solid (1.38 g, 82%).**^1^H NMR** (DMSO-d_6_): *δ* 10.45 (s, 1H), 8.75 (s, 1H), 7.06 (s, 2H), 3.85 (s, 6H), 3.68 (s, 3H). **^13^C NMR** (DMSO-d_6_): *δ* 160.9, 158.5, 158.0, 153.2, 134.8, 134.5, 124.8, 122.1, 11.4, 99.2, 60.7, 56.3. LC-MS (ESI) calcd. for C_14_H_13_ClF_3_N_3_O_3_ [M+H]^+^: 364.05; found: 363.95.

**4-Chloro-*N*-(5-methoxy-2-methylphenyl)-5-(trifluoromethyl)pyrimidin-2-amine (5c).** The title compound was prepared by the reaction of 5-trifluromethyl-2,4-dichloropyrimidine (1 g, 4.6 mmol), ZnCl_2_ (5.5 mL, 5.5 mmol), 5-methoxy-2-methylaniline (0.60 g, 4.6 mmol) and triethylamine (1 g, 5.5 mmol) according to general method D. Colorless solid (0.800 g, 53%).**^1^H NMR** (DMSO-d_6_): *δ* 8.04 (s, 1H), 7.20 (d, *J* = 2.8 Hz, 1H), 7.19 (d, *J* = 8.2 Hz, 1H), 6.68 (dd, *J* = 8.9 Hz, 2.8 Hz, 1H), 3.70 (s, 3H), 2.11 (s, 3H). **^13^C NMR** (DMSO-d_6_): *δ* 158.9, 158.2, 155.6, 154.3, 135.9, 131.5, 124.3, 118.8, 111.6, 55.7, 17.5. LC-MS (ESI) calcd. for C_13_H_11_ClF_3_N_2_O [M+H]^+^: 318.05; found: 318.00.

**4-Chloro-*N*-(2-methoxy-4-morpholinophenyl)-5-(trifluoromethyl)pyrimidin-2-amine (5d).** The title compound was prepared by the reaction of 5-trifluromethyl-2,4-dichloropyrimidine (1 g, 4.6 mmol), ZnCl_2_ (5.5 mL, 5.5 mmol), 2-methoxy-4-morpholinoaniline (0.952 g, 4.6 mmol) and triethylamine (1 g, 5.5 mmol) according to general method D. Yellow solid (0.784 g, 43.6%). ^1^H-NMR (CDCl_3_) δ: 8.53 (s, 1H), 8.15 (d, *J* = 8.2 Hz, 1H), 7.86 (brs, 1H), 6.56-6.52 (m 2H), 3.89-3.85 (overlapping singlet and triplet, 7H), 3.14 (t, *J* = 4.6 Hz, 4H). LC-MS (ESI) calcd. for C_16_H_16_ClF_3_N_4_O_2_ [M+H]^+^: 389.09; found: 388.90.

**2-(5-Chloro-2-(2-methoxy-4-morpholinophenylamino)pyrimidin-4-ylamino)-N-methylbenzamide (6a)**. 2-(2,5-Dichloropyrimidin-4-ylamino)-*N*-methylbenzamide (**3a**, 0.148 g, 0.5 mmol), 2-methoxy-4-morpholinoaniline (0.208 g, 1 mmol) and HCl were processed according to General method A. Brown solid (0.150, 64%). **^1^H NMR** (DMSO-d_6_): 11.56 (s, 1H), 8.68 (d, *J* = 4.6 Hz, 2H), 8.56 (d, *J* = 7.8 Hz, 1H), 8.14 (s, 1H), 7.68 (d, *J* = 9.2 Hz, 1H), 7.39 (d, *J* = 8.7 Hz, 1H), 7.29 (t, *J* = 7.8 Hz, 1H), 7.03 (t, *J* = 15.0 Hz, 1H), 6.61 (d, *J* = 2.3 Hz, 1H), 6.46 (dd, *J* = 11.5, 2.8 Hz, 1H), 3.73-3.64 (overlapping singlet and triplet, 7H), 3.09 (t, *J* = 4.6 Hz, 4H), 2.75 (d, *J* = 4.6 Hz, 3H). **^13^C NMR** (DMSO-d_6_): *δ* 169.5, 164.0, 159.5, 155.4, 155.2, 149.7, 140.1, 131.9, 128.4, 129.9, 122.1, 121.5, 120.9, 120.6, 106.9, 104.5, 100.3, 66.7, 55.6, 49.6, 26.8. LC-MS (ESI) calcd. for C_23_H_25_ClN_6_O_6_[M+H]^+^: 469.17; found: 469.10. HRMS (ESI) Calcd for C_23_H_25_ClN_6_O_6_[M+H]^+^: 469.1749; Found: 469.1749.

**2-(5-Bromo-2-(2-methoxy-4-morpholinophenylamino)pyrimidin-4-ylamino)-*N*-methylbenzamide (6b).** 2-(5-Bromo-2-chloropyrimidin-4-ylamino)-*N*-methylbenzamide (**3b**, 0.341 g, 1 mmol), 2-methoxy-4-morpholinoaniline (0.208 g, 1 mmol) and HCl were processed according to general method C. Tan solid (0.258 g, 50%). **^1^H NMR** (DMSO-d_6_): *δ* 11.33 (s, 1H), 8.80 (brs, 1H), 8.50 (brs, 1H), 8.01(s, 1H), 7.64 (d, *J* = 7.3 Hz, 1H), 7.34 (d, *J* = 8.2 Hz, 1H), 7.27 (t, *J* = 7.2 Hz, 1H), 7.03 (t, *J* = 7.3 Hz, 1H), 6.61 (s, 1H), 6.43 (d, *J*= 8.7 Hz, 1H), 5.71 (s, 1H), 3.72-3.68 (overlapping singlet and triplet, 7H), 3.06 (t, *J* = 4.6 Hz, 4H), 2.75 (d, *J* = 4.3 Hz, 3H). **^13^C NMR** (DMSO-d_6_): *δ* 169.4, 159.9, 158.1, 156.2, 153.3, 149.7, 140.0, 131.8, 128.3, 125.9, 122.1, 121.7, 121.0, 120.8, 106.9, 100.4, 93.7, 66.7, 55.9, 49.6, 26.8. LC-MS (ESI) calcd. for C_23_H_25_BrN_6_O_3_ [M+H]^+^: 515.12; found: 515.05. HRMS (ESI) calcd. for C_23_H_25_BrN_6_O_3_ [M+H]^+^: 515.1227; found: 515.1226.

**2-(2-(2-Methoxy-4-morpholinophenylamino)-5-(trifluoromethyl)pyrimidin-4-ylamino)-*N*-methylbenzamide (6c).** 4-Chloro-*N*-(2-methoxy-4-morpholinophenyl)-5-(trifluoromethyl)pyrimidin-2-amine (**5d**, 0.194 g, 0.5 mmol), 2-amino-*N*-methylbenzamide and DIEA (0.075 g, 0.5 mmol) were processed according to general method E. to afford the title compound as a colorless solid (0.125 g, 50%). **^1^H NMR** (DMSO-d_6_): *δ* 11.40 (s, 1H), 8.65 (s, 1H), 8.64 (s, 2H), 8.27 (s, 1H), 7.62 (d, *J* = 7.8 Hz, 1H), 7.26-7.24 (m, 2H), 7.03 (t, *J* = 7.7 Hz, 1H), 6.62 (d, *J* = 2.3 Hz, 1H), 6.45 (dd, *J*_1_ = 10.0 Hz, *J*_2_ = 2.8 Hz, 1H), 3.73-3.71 (overlapping singlet and triplet, 7H), 3.09 (t, *J* = 5.2 Hz, 4H), 2.73 (d, *J* = 4.6 Hz, 3H). **^13^CNMR** (DMSO-d_6_): *δ* 169.4, 162.7, 156.5, 151.0, 139.6, 131.7, 128.2, 122.8, 122.4, 121.5, 119.7, 106.9, 100.2, 66.7, 58.0, 49.9, 26.7. LC-MS (ESI) calcd. for C_24_H_25_F_3_N_6_O_3_ [M+H]^+^: 503.19; found: 503.05. HRMS (ESI) calcd. for C_24_H_25_F_3_N_6_O_3_ [M+H]^+^: 503.2013; found: 503.2001.

***N*-Methyl-2-(2-(2-oxo-1,2,3,4-tetrahydroquinolin-6-ylamino)-5-(trifluoromethyl)pyrimidin-4-ylamino)benzamide (6d).** 6-(4-Chloro-5-(trifluoromethyl)pyrimidin-2-ylamino)-3,4-dihydroquinolin-2(1H)-one (**5a**, 0.085, 0.25 mmol), 2-amino-*N*-methyl benzamide (0.37 g, 0.25 mmol) and DIEA (0.035 g, 0.25 mmol) were processed according to general method E. to afford the title compound as a colorless solid (0.085 g, 75%). **^1^H NMR** (DMSO-d_6_): *δ* 11.32 (s, 1H), 9.95 (s, 1H), 9.73 (s, 1H), 8.70 (s, 1H), 8.37 (s, 1H), 7.67 (d, *J* = 7.8 Hz, 1H), 7.52-7.23 (m, 4H), 7.12 (t, *J* = 7.8 Hz, 1H)), 6.72 (d, *J* = 8.7 Hz, 1H), 2.74-2.73 (overlapping doublets and triplets, 5H), 2.46 (t, *J* = 7.3 H, 2H). **^13^C NMR** (DMSO-d_6_): *δ* 170.4, 169.3, 160.9, 158.9, 155.8, 143.9, 134.3, 133.9, 132.0, 128.4, 124.2, 123.1, 121.0, 120.4, 115.4, 30.9, 26.7, 25.6. LC-MS (ESI) calcd. for C_22_H_19_F_3_N_6_O_2_ [M+H]^+^: 457.15; found: 457.05. HRMS (ESI) calcd. for C_22_H_19_F_3_N_6_O_2_ [M+H]^+^: 457.1594; found: 457.1585.

**2-(5-Chloro-2-(2-oxo-1,2,3,4-tetrahydroquinolin-6-ylamino)pyrimidin-4-ylamino)-*N*-methylbenzamide (6e).** 2-(2,5-Dichloropyrimidin-4-ylamino)-*N*-methylbenzamide (**3a**, 0.148 g, 0.5 mmol) and 6-amino-3,4-dihydroquinolin-2(1H)-one (0.162 g, 1 mmol) and DIEA were processed according to general method B. to afford the desired compound as a tan solid (0.154 g, 73%). **^1^H NMR** (DMSO-d_6_): *δ* 11.87 (s, 1H), 9.99 (s, 1H), 9.74 (s, 1H), 8.80 (d, *J* = 4.1 Hz, 1H), 8.56 (brs, 1H), 8.22 (s, 1H), 7.74 (d, *J* = 7.3 Hz, 1H), 7.40-7.38 (m, 2H), 7.24 (d, *J* = 7.8 Hz, 1H), 7.15 (t, *J* = 7.3 Hz, 1H), 6 77 (d, *J* = 8.7 Hz, 1H), 2.72-2.76 (overlapping doublet and triplet, 5H), 2.39 (t, *J* = 7.3 Hz, 2H). **^13^C NMR** (DMSO-d_6_): *δ* 170.8, 169.4, 155.9, 155.1, 138.2, 134.8, 133.5, 132.0, 128.5, 124.3, 123.4, 122.5, 121.9, 121.4, 119.8, 117.4, 109.3, 30.9, 26.9, 25.6. LC-MS (ESI) calcd. for C_21_H_19_ClN_6_O_2_ [M+H]^+^: 423.13; found: 423.00. HRMS (ESI) calcd. for C_21_H_19_ClN_6_O_2_ [M+H]^+^: 423.1331; found: 423.1303.

**2-(5-Chloro-2-(2-oxoindolin-5-ylamino)pyrimidin-4-ylamino)-*N*-methylbenzamide (6f)**. **2-**(2,5-Dichloropyrimidin-4-ylamino)-*N*-methylbenzamide (**3a**, 0.148 g, 0.5 mmol), 5-aminoindolin-2-one (0.148 g, 1 mmol) were taken in ^n^BuOH. It was then processed according to general method B. to afford the desired compound as a tan solid (0.147 g, 73%). **^1^H NMR** (DMSO-d_6_): *δ* 11.50 (s, 1H), 10.23 (s, 1H), 9.29 (s, 1H), 8.71 –8.70 (overlapping singlet and doubletet, 2H), 8.13 (s, 1H), 7.70 (d, *J* = 7.8 Hz, 1H), 7.56 (s, 1H), 7.43 (t, *J* = 7.3 Hz, 1H), 7. 30 (d, *J* = 7.8 Hz, 1H), 7.10 (t, *J* = 7.3 Hz, 1H), 6.70 (d, *J* = 8.2 Hz, 1H), 3.39 (s, 2H), 2.76 (d, *J* = 3.2 Hz, 3H). **^13^CNMR** (DMSO-d_6_): *δ* 176.8, 169.4, 158.4, 155.5, 155.1, 139.8, 138.9, 134.8, 132.0, 128.5, 126.4, 122.5, 121.9, 121.4, 119.8, 117.4, 109.3, 35.6, 26.8. LC-MS (ESI) calcd. for C_20_H_17_ClN_6_O_2_ [M+H]^+^: 409.11; found: 409.05.

**2-(2-(1H-indol-5-ylamino)-5-chloropyrimidin-4-ylamino)-*N*-methylbenzamide (6g).** The title compound was prepared from 2-(2,5-dichloropyrimidin-4-ylamino)-*N*-methylbenzamide (**3a**, 0.148 g, 0.5 mmol) and 1H-indol-5-amine (0.132 g, 1 mmol) were processed according to general method B. Tan solid (0.172 g, 84%). **^1^H NMR** (DMSO-d_6_): *δ* 11.03 (s, 1H), 9.66 (s, 1H), 8.63-8.51 (m, 1H), 7.85 (s, 1H), 7.70 (s, 1H), 7.53-7.38 (m, 1H), 7.14 (d, *J* = 7.8 Hz, 1H), 7.24 (s, 1H), 7.15-7.09 (m, 3H), 7.00-6.96 (m, 1H), 6.81 (t, *J* = 7.8 Hz, 1H), 6.21 (s, 1H), 2.77 (d, *J* = 4.2 Hz, 3H). **^13^C NMR** (DMSO-d_6_): *δ* 169.7, 158.7, 155.6, 154.3, 139.7, 131.6, 131.5, 128.0, 127.5, 125.2, 122.1, 121.6, 121.5, 117.2, 112.9, 111.2, 101.6, 26.7. LC-MS (ESI) calcd. for C_20_H_17_ClN_6_O [M+H]^+^: 393.12; found: 392.95. HRMS (ESI) calcd. for C_20_H_17_ClN_6_O [M+H]^+^: 393.1152; found: 393.1213.

**2-(2-(1H-indol-5-ylamino)-5-bromopyrimidin-4-ylamino)-*N*-methylbenzamide (6h).** 2-(5-Bromo-2-chloropyrimidin-4-ylamino)-*N*-methylbenzamide (**3b**, 0.341 g, 1 mmol) 1H-indol-5-amine (0.264 g, 2 mmol) were processed according to general method B. to afford the desired compound as a tan solid (0.296 g, 68%). **^1^H NMR** (DMSO-d_6_): *δ* 11.28 (s, 1H), 10.91 (s, 1H), 9.18 (s, 1H), 8.68-8.67 (m, 2H), 8.19 (s, 1H), 7.83 (s, 1H), 7.66 (d, *J* = 7.8 Hz, 1H), 7.27-7.18 (m, 4H), 7.06 (t, *J* = 7.3 Hz, 1H), 6.29 (s, 1H), 2.76 (d, *J* = 4.6 Hz, 3H). **^13^CNMR** (DMSO-d_6_): *δ* 169.4, 159.4, 158.1, 156.2, 139.9, 132.9, 132.4, 131.8, 128.4, 128.1, 126.2, 122.2, 117.1, 111.4, 101.4, 26.9. LC-MS (ESI) calcd. for C_20_H_17_BrN_6_O [M+H]^+^: 439.06; found: 439.00. HRMS (ESI) calcd. for C_20_H_17_BrN_6_O [M+H]^+^: 439.0702; found: 439.0693.

**2-(5-Chloro-2-(6-methylbenzo[d][1,3]dioxol-5-ylamino)pyrimidin-4-ylamino)-N-methylbenzamide (6i)**. 2-(5-Chloro-2-chloropyrimidin-4-ylamino)-*N*-methylbenzamide (**3a**, 0.296 g, 1 mmol) and 6-methylbenzo[d][1,3]dioxol-5-amine (0.151 g, 1 mmol) were processed according to general method B. brown solid (0.286 g, 69%). **^1^H NMR** (DMSO-d_6_): *δ* 11.62 (s, 1H), 8.68-8.67 (m, 1H), 8.62 (s, 1H), 8.52 (d, *J* = 6.4 Hz, 1H), 8.05 (s, 1H), 7.65 (dd, *J* = 7.8 Hz, 1.8 Hz, 1H), 7.20 (t, *J* = 8.7 Hz, 1H), 7.02 (t, *J* = 7.8 Hz, 1H), 6.89 (s, 1H), 6.79 (s, 1H), 5.90 (s, 2H), 2.75 (d, *J* = 4.6 Hz, 3H), 2.10 (s, 3H). **^13^C NMR** (DMSO-d_6_): *δ* 169.5, 159.9, 155.5, 155.4, 140.2, 128.4, 127.2, 122.0, 121.2, 120.4, 110.0, 108.6, 104.5, 101.4, 72.9, 68.4, 60.7, 26.8, 18.4. LC-MS (ESI) calcd. for C_20_H_18_ClN_5_O_3_[M+H]^+^: 412.11; found: 412.00. HRMS (ESI) calcd. for C_20_H_18_ClN_5_O_3_ [M+H]^+^: 412.1171; found: 413.1158.

**2-(5-Chloro-2-(2,3-dihydrobenzo[b][1,4]dioxin-6-ylamino)pyrimidin-4-ylamino)-*N*-methylbenzamide (6j).** 2-(2,5-Dichloropyrimidin-4-ylamino)-*N*-methylbenzamide (**3a**, 0.148 g, 0.5 mmol) and 2,3-dihydrobenzo[b][1,4]dioxin-6-amine (0.151g, 1 mmol) were taken in ^n^BuOH. It was then processed according to general method B. to afford the desired compound as a tan solid (0.172 g, 84%). **^1^H NMR** (DMSO-d_6_): *δ* 11.56 (s, 1H), 9.22 (s, 1H), 8.71-8.70 (m, 2H), 8.14 (s, 1H), 7.70 (d, *J* = 7.3 Hz, 1H), 7.43 (t, *J* = 7.8 Hz, 1H), 7.23 (s, 1H), 7.08 (t, *J* = 7.3 Hz, 1H), 7.00 (s, 1H), 6.71 (d, *J* = 8.7 Hz, 1H), 4.18 (t, *J* = 5.5 Hz, 4H), 2.77 (d, *J* = 4.6 Hz, 3H). **^13^CNMR** (DMSO-d_6_): *δ* 169.5, 158.3, 155.4, 155.1, 143.4, 139.9, 138.9, 134.4, 132.1, 128.5, 122.4, 121.7, 121.1, 116.9, 113.7, 109.5, 64.7, 64.5, 26.8. LC-MS (ESI) calcd. for C_20_H_18_ClN_5_O_3_[M+H]^+^: 412.11; found: 412.00. HRMS (ESI) calcd. for C_20_H_18_ClN_5_O_3_[M+H]^+^: 412.1171; found: 412.1151.

**2-(5-Chloro-2-(5-methoxy-2-methylphenylamino)pyrimidin-4-ylamino)-*N*-methylbenzamide (6k).** 2-(2,5-Dichloropyrimidin-4-ylamino)-*N*-methylbenzamide (**3a**, 0.296 g, 1 mmol) and 5-methoxy-2-methylaniline (0.274 g, 2 mmol) were processed according to general method B. Yellow solid (0.232 g, 58%). **^1^H NMR** (DMSO-d_6_): *δ* 11.44 (s, 1H), 8.48 (s, 1H), 8.66 (d, *J* = 8.7 Hz, 1H), (d, *J* = 8.7 Hz, 1H), 8.16 (s, 1H), 7.64 (d, *J* = 7.8 Hz,1H), 7.09-7.04 (m, 4H), 6.65 (d, *J* = 8.2 Hz,1H), 3.65 (s, 3H), 2.77 (d, *J* = 4.6 Hz, 3H), 2.11 (s, 3H). **^13^C NMR** (DMSO-d_6_): *δ* 169.4, 159.5, 158.3, 158.0, 156.7, 140.1, 139.2, 131.7, 131.2, 128.4, 125.1, 122.1, 121.4, 112.2, 110.7, 94.0, 55.7, 26.8, 17.7. LC-MS (ESI) calcd. for C_20_H_20_ClN_5_O_2_ [M+H]^+^: 398.13; found: 398.00. HRMS (ESI) calcd. for C_20_H_20_ClN_5_O_2_ [M+H]^+^: 398.1378; found: 398.1366.

**2-(2-(5-Methoxy-2-methylphenylamino)-5-(trifluoromethyl)pyrimidin-4-ylamino)-*N*-methylbenzamide (6l).** 4-Chloro-*N*-(5-methoxy-2-methylphenyl)-5-(trifluoromethyl)pyrimidin-2-amine (**5c**, 0.158 g, 0.5 mmol), 2-amino-*N*-methylbenzamide (0.075 g, 0.5 mmol) and DIEA (0.066 g, 0.5 mmol) were processed according to general method E. to afford the title compound as a colorless solid (0.186 g, 86%). **^1^H NMR** (DMSO-d_6_): *δ* 11.53 (s, 1H), 9.26 (s, 2H), 8.67-8.66 (m, 2H), 8.32 (s, 1H), 7.63(dd, *J* = 7.8 Hz, 1.7 Hz, 1H), 7.11 (d, *J* = 8.2 Hz, 1H), 7.01 (t, *J* = 7.3 Hz, 1H), 6.92 (d, *J* = 2.3 Hz, 1H), 6.73(dd, *J=* 8.2 Hz, 2.8 Hz, 1H), 3.62 (s, 3H), 2.71 (d, *J* = 4.6 Hz, 3H), 2.05 (s, 3H). **^13^C NMR** (DMSO-d_6_): *δ* 169.4, 162.3, 158.1, 156.5, 156.2, 139.5, 138.3, 131.5, 131.3, 128.2, 125.9, 122.6, 122.5, 121.3, 113.0, 111.8, 55.6, 26.8, 17.6. LC-MS (ESI) calcd. for C_21_H_20_F_3_N_5_O_2_ [M+H]^+^: 432.15; found: 432.05. HRMS (ESI) calcd. for C_21_H_20_F_3_N_5_O_2_ [M+H]^+^: 432.1642; found: 432.1631.

**2-(5-Bromo-2-(5-methoxy-2-methylphenylamino)pyrimidin-4-ylamino)-*N*-methylbenzamide (6m).** 2-(5-Bromo-2-chloropyrimidin-4-ylamino)-*N*-methylbenzamide (**3b**, 0.341 g, 1 mmol) and 5-methoxy-2-methylaniline (0.274 g, 2 mmol) were processed according to general method B. Tan solid (0.298 g, 67%). **^1^H NMR** (DMSO-d_6_): *δ* 11.61 (s, 1H), 9.40 (s, 1H), 8.73-8.72 (m, 2H), 8.20 (s, 1H), 7.72 (d, *J* = 7.8 Hz, 1H), 7.44 (t, *J* = 7.4 Hz, 1H), 7.30-7.10 (m, 3H), 6.51 (d, *J* = 7.8 Hz, 1H), 3.65 (s, 3H), 2.77 (d, *J* = 4.1 Hz, 3H), 2.11 (s, 3H). **^13^C NMR** (DMSO-d_6_): *δ* 168.9, 159.5, 158.2, 155.5, 155.1, 141.9, 139.8, 132.1, 129.6, 129.1, 111.9, 107.4, 106.5, 105.6, 54.9, 26.3. LC-MS (ESI) calcd. for C_20_H_20_BrN_5_O_2_[M+H]^+^: 444.08; found: 443.95. HRMS (ESI) calcd. for C_20_H_20_BrN_5_O_2_[M+H]^+^: 444.0855; found: 444.0849.

**2-(5-Chloro-2-(3-methoxyphenylamino)pyrimidin-4-ylamino)-*N*-methylbenzamide (6n)**. **2-**(2,5-Dichloropyrimidin-4-ylamino)-*N*-methylbenzamide (**3a**, 0.297 g, 1 mmol) and 3-methoxyaniline (0.246 g, 2 mmol) were taken in ^n^BuOH. It was then processed according to general method B to afford the desired compound as a colorless solid (0.253 g, 66%). **^1^H NMR** (DMSO-d_6_): *δ* 11.6 (s, 1H), 9.40 (s, 1H), 8.73-8.72 (m, 2H), 8.19 (s, 1H), 7.29 (d, *J* = 8.8 Hz, 1H), 7.14 (t, *J* = 7.3 Hz, 1H), 7.12-7.10 (m, 4H), 6.51 (d, *J* = 7.8 Hz, 1H), 3.65 (s, 3H), 2.77 (d, *J* = 4.1 Hz, 3H). **^13^C NMR** (DMSO-d_6_): *δ* 168.9, 159.5, 158.0, 152.8, 152.0, 141.4, 139.8, 132.1, 129.6, 129.0, 122.4, 121.8, 121.1, 11.9, 107.4, 106.6, 105.5, 54.9, 26.3. LC-MS (ESI) calcd. for C_19_H_18_ClN_5_O_2_ [M+H]^+^: 384.11; found: 384.00. HRMS (ESI) calcd. for C_19_H_18_ClN_5_O_2_ [M+H]^+^: 384.1222; found: 384.1210

**2-(5-Chloro-2-(3,4,5-trimethoxyphenylamino)pyrimidin-4-ylamino)-*N*-methylbenzamide (6o).** A mixture of 2-(2,5-dichloropyrimidin-4-ylamino)-*N*-methylbenzamide (**3a**, 0.296 g, 1 mmol) and 3,4,5-trimethoxy aniline (0.366 g, 2 mmol) were processed according to general method B. Colorless solid (0.160, 73%). **^1^H NMR** (DMSO-d_6_): *δ* 11.63 (s, 1H), 9.27 (s, 1H), 8.27-8.71 (m, 2H), 8.18 (s, 1H), 7.70 (d, *J* = 9.2 Hz, 1H), 7.69 (t, *J* = 7.8 Hz, 1H), 7.08 (t, *J* = 7.3 Hz, 1H), 6.98 (s, 2H), 3.62 (s, 3H), 3.59 (s, 6H), 2.76 (d, *J* = 4.6 Hz, 3H). **^13^C NMR** (DMSO-d_6_): *δ* 169.5, 158.3, 155.4, 155.1, 153.2, 139.9, 136.8, 133.2, 132.0, 128.6, 122.3, 121.4, 120.9, 105.7, 98.5, 60.7, 56.2, 26.8. LC-MS (ESI) calcd. for C_21_H_22_ClN_5_O_4_ [M+H]^+^: 444.13; found: 444.05. HRMS (ESI) calcd. for C_21_H_22_ClN_5_O_4_ [M+H]^+^: 444.1433; found: 444.1431.

***N*-Methyl-2-(5-(trifluoromethyl)-2-(3,4,5-trimethoxyphenylamino)pyrimidin-4-ylamino)benzamide (6p).** 4-Chloro-5-(trifluoromethyl)-*N*-(3,4,5-trimethoxyphenyl)pyrimidin-2-amine (**5b**, 0.363 g, 1 mmol), 2-amino-*N*-methylbenzamide and DIEA (0.150 g, 1 mmol) were processed according to general method E. to afford the title compound as a colorless solid (0.232 g, 59%). **^1^H NMR** (DMSO-d_6_): *δ* 11.43 (s, 1H), 9.65 (s, 1H), 8.72 (d, *J* = 4.0 Hz, 1H), 8.42-8.40 (2H), 7.67 (d, *J* = 7.8 Hz, 1H), 7.31 (s, 1H), 7.10 (t, *J* = 7.3 Hz, 1H), 6.95 (s, 2H), 3.68 (s, 6H), 3.58 (s, 3H), 2.75 (d, *J* = 4.6 Hz, 3H). **^13^C NMR** (DMSO-d_6_): 169.4, 161.3, 156.4, 153.1, 139.4, 135.9, 133.9, 131.8, 128.4, 126.4, 123.7, 122.9, 121.9, 99.5, 60.6, 56.2, 26.8. LC-MS (ESI) calcd. for C_22_H_22_F_3_N_5_O_4_ [M+H]^+^: 478.16; found: 478.10. HRMS (ESI) calcd. for C_22_H_22_F_3_N_5_O_4_ [M+H]^+^: 478.1683; found: 478.1683.

***N*-Methyl-2-(2-(3,4,5-trimethoxyphenylamino)pyrimidin-4-ylamino)benzamide (6q).** A mixture of 2-amino-*N*-methyl benzamide (0.015 mg, 0.1 mmol), 2,4-dichloro pyrimidine (0.015 mg, 0.1 mmol) and DIEA (0.013 g, 0.1 mmol) in ^n^BuOH. The reaction mixture was stirred at 110°C for 4-5 h. The reaction mixture was cooled to room temperature and 3,4,5-trimethoxyaniline (0.018 mg, 0.1 mmol) and DIEA (0.013 g, 0.1 mmol) were added and heated at 110 °C for 18 h. The reaction mixture was stirred at 110 °C for 20 h. It was then cooled to room temperature excess solvent was reduced under reduced pressure. The crude residue was purified using automated prep-HPLC to yield the title compound. Colorless solid (0.036 g, 88%). **^1^H NMR** (DMSO-d_6_): *δ* 11.55 (s, 1H), 10.11 (brs, 1H), 8.55 (d, *J* = 4.6 Hz, 1H), 8.08 (d, *J* = 7.8 Hz, 1H), 7.93 (d, *J* = 6.5 Hz, 1H), 7.63 (d, *J* = 7.8 Hz, 1H), 7.36 (t, *J* = 7.8 Hz, 1H), 7.20 (t, *J* = 8.6 Hz, 1H), 6.89 (s, 2H), 6.42 (d, *J* = 6.5 Hz, 1H),3.63 (s, 3H), 3.61 (s, 9H), 2.73 (d, *J* = 4.6 Hz, 3H). **^13^C NMR** (DMSO-d_6_): *δ* 168.6, 165.0, 161.3, 156.6, 138.0, 135.3, 131.5, 128.8, 125.5, 124.0, 101.9, 100.9, 66.6, 56.3, 27.3. LC-MS (ESI) calcd. for C_21_H_23_N_5_O_4_ [M+H]^+^:410.18; found: 410.05. HRMS (ESI) calcd. for C_21_H_23_N_5_O_4_ [M+H]^+^:410.1823; found: 410.1813.

**2-(2-(3,5-dimorpholinophenylamino)-5-(trifluoromethyl)pyrimidin-4-ylamino)-*N*-methylbenzamide (6r)**. 4-Chloro-N-(3,5-dimorpholinophenyl)-5-(trifluoromethyl)pyrimidin-2-amine (0.110 g, 0.25 mmol), and 2-amino-*N*-methyl benzamide (0.041 g, 0.275 mmol) and HCl were processed according to general method C to afford the title compound as a brown solid (0.097 g, 70%). **^1^H NMR** (DMSO-d_6_): *δ* 11.48 (s, 1H), 9.60 (s, 1H), 8.71 (s, 1H), 8.40 (s, 1H), 7.69 (d, *J* = 7.8 Hz, 2H), 7.36 (s, 1H), 7.10 (t, *J* = 7.3 Hz, 1H), 6.77 (s, 1H), 6.24 (s, 2H), 3.63 (brs, 8H), 2.96 (brs, 8H), 2.75 (d, *J* = 4.7 Hz, 3H). **^13^C NMR** (DMSO-d_6_): *δ* 169.4, 161.1, 158.9, 157.6, 156.4, 156.0, 152.0, 141.0, 139.0, 132.0, 128.3, 122.8, 122.7, 121.8, 100.8, 98.9, 66.6, 49.4, 26.8. LC-MS (ESI) calcd. for C_27_H_30_F_3_N_7_O_3_ [M+H]^+^: 558.24; found: 558.20. HRMS (ESI) calcd. for C_27_H_30_F_3_N_7_O_3_ [M+H]^+^: 558.2435; found: 558.2424.

**2-(5-Chloro-2-(3,5-dimorpholinophenylamino)pyrimidin-4-ylamino)-N-methylbenzamide (6s).** 2-(2,5-Dichloropyrimidin-4-ylamino)-*N*-methylbenzamide (**3a,** 0.148 g, 0.5 mmol) and 3,5-dimorpholinoaniline (0.263 g, 1 mmol) were processed according to general method B to afford the title compound as a brownish solid (0.170 g, 65%). **^1^H NMR** (DMSO-d_6_): *δ* 11.77 (s, 1H), 9.35 (s, 1H), 8.77-8.76 (m, 2H), 8.22 (s, 1H), 7.72 (dd, *J* = 8.2 Hz, 1.3 Hz, 1H), 7.41 (t, *J* = 7.3 Hz, 1H), 7.10 (t, *J* = 7.8 Hz, 1H), 6.78 (s, 2H), 6.23 (s, 1H), 3.71 (t, *J* = 4.6 Hz, 8H),. 3.01 (t, *J* = 5.0 Hz, 8 H), 2.77 (d, *J* = 4.6 Hz, 3H). **^13^C NMR** (DMSO-d_6_): *δ* 169.5, 158.8, 155.5, 153.2, 142.0, 139.8, 132.3, 128.8, 121.4, 121.0, 105.2, 100.4, 98.0, 66.5, 49.5, 26.8. LC-MS (ESI) calcd. for C_26_H_30_ClN_7_O_3_ [M+H]^+^: 524.20; found: 524.20. HRMS (ESI) calcd. for C_26_H_30_ClN_7_O_3_ [M+H]^+^: 524.2171; found: 524.2158.

**2-(5-Bromo-2-(3,5-dimorpholinophenylamino)pyrimidin-4-ylamino)-N-methylbenzamide (6t).** 2-(5-Bromo-2-chloropyrimidin-4-ylamino)-*N*-methylbenzamide (**3b**, 0.171 g, 0.5 mmol) and 3,5-dimorpholinoaniline (0.263 g, 1 mmol) were processed according to general method B. to afford the title compound as a tan solid (0.177 g, 62%). **^1^H NMR** (DMSO-d_6_): *δ* 11.37 (s, 1H), 9.13 (s, 1H), 8.71-8.62 (m, 2H), 8.24 (s, 1H), 7.68 (dd, *J* = 9.16 Hz, 1.3 Hz, 1H), 7.41 (t, *J* = 7.3 Hz, 1H), 7.10 (t, *J* = 7.8 Hz, 1H), 6.73(s, 2H), 6.11 (s, 1H), 3.62 (t, *J* = 4.1 Hz, 8H), 2.94 (t, *J* = 4.6 Hz, 8 H), 2.77 (d, *J* = 4.6 Hz, 3H). **^13^C NMR** (DMSO-d_6_): *δ* 168.9, 1582, 157.5, 155.6, 153.2,152.1, 141.2, 139.3, 131.6, 127.9, 121.8, 121.1, 120.7, 99.1, 97.3, 94.1, 66.2, 48.8, 26.3. LC-MS (ESI) calcd. for C_26_H_30_BrN_7_O_3_ [M+H]^+^: 568.48; found: 568.50.

### Cell Culture

Kras^LSL-G12D/+^;Trp53^LSL-R172H/+^;Pdx1-Cre (KPC) mice-derived PDAC cells (KPC.4662) were provided by Dr. Robert Vonderheide (University of Pennsylvania) and cultured in high glucose, L-glutamine DMEM supplemented with 10% FBS, 100 U/mL penicillin-streptomycin (pen-strep), and 20 mM HEPES. Other cell lines (A549 – CRM-CCL-185, H69AR – CRL-11351, MiaPaca2 – CRM-CRL-1420, BxPC3 – CRL-1687, Panc1 – CRL-1469, and NCI-H358 – CRL-5807) were purchased via ATCC and cultured in recommended media.

### shRNA Knockdown

For the viral infection, cells were plated at 1×10^5^/well of 6-well plates and incubated for 24 h. The viruses were mixed with 8 μg/mL Polybrene (Santa Cruz Biotechnology #sc-134220) and added to the cells. Three days later, infected cells were selected by the addition of 1 μg/mL puromycin (Gibco #A11138-03) for an additional 3 days, harvested by trypsinization, and dispensed into 96-well plates for experiments._After knockdown of the autophagy genes, cell survival and proliferation indicated by cell viability over 8 days was measured with CellTiter-Glo and plotted. Compared with control cells, knockdown of ULK1 or ATG13 significantly reduced the proliferation of all three cell lines. After knockdown of the autophagy genes, cell survival and proliferation indicated by cell viability over 8 days was measured with CellTiter-Glo and plotted.

### shRNA target sequence for the knockdown study

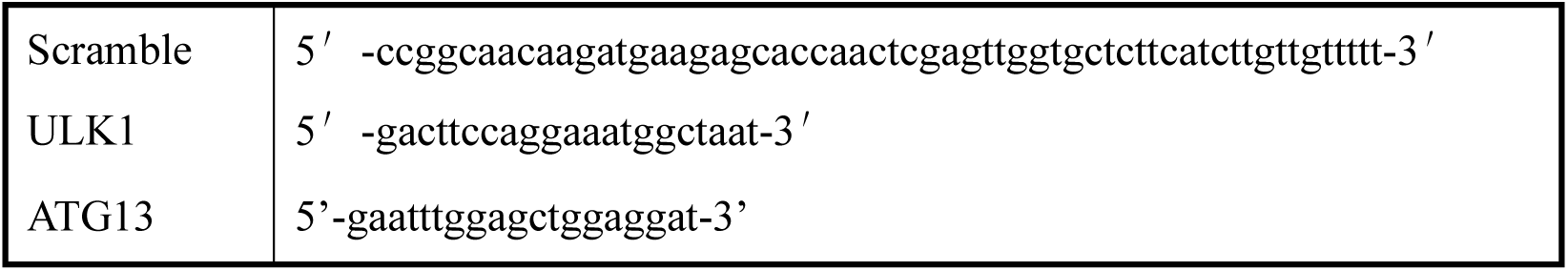

### CRISPR Knockouts for Autophagy Genes

Small Guide RNAs (sgRNAs) targeting human ATG7, ATG13, FIP200, and ULK1 (guide JG) were described previously ^52^. sgRNAs targeting ATG5, ULK1 (g1B), ULK2 (gA and gC) were selected using the optimized CRISPR design tool (http://crispr.mit.edu). Guides with high targeting scores and low probability of off-target effects were chosen, targeting the 1st or 2nd coding exon, or otherwise the first most common downstream exon for transcripts reported to produce multiple isoforms from searches of Uniprot or Ensembl databases. Non-targeting guide was previously described ^53^. Oligonucleotide sequences are listed in **Supplemental Table S1**. Oligonucleotides for sgRNAs were synthesized by IDT, annealed in vitro and subcloned into BsmbI-digested lentiCRISPRv.2-puro (Addgene 52961) or lentiCas9-Blast (ULK2 sgRNAs, Addgene 52962). Stable CRISPR knockout pools of MiaPaca2 or BxPC3 were generated by stable integration of pLentiCRISPRv.2 and lentiCas9-Blast plasmids by lentiviral transduction with 0.45um-filtered viral supernatant supplemented with polybrene followed by selection with 2ug/ml puromycin and 20ug/ml blasticidin. Validation of guide specificity and degree of knockout was assessed by Western blot of low-passage MiaPaca2 or BxPC3 cell pools and low-passage cells KO lines were used for Cyquant, low density, and soft agar assays. Polyclonal cell pools were injected in flanks of nude mice for tumor growth studies.

### In Vivo ULK CRISPR Knockout

All procedures using animals were approved by the Salk Institute Institutional Animal Care and Use Committee (IACUC). For flank tumor studies, 6-8 week old female J;Nu mice were purchased from Jackson labs (007850). 5 million MiaPaca2 cells stably expressing sgRNAs targeting ULK1 (g1B) or ULK2 (g2C) or non-targeting control in 100ul of 1:1 PBS and Matrigel (BD 354234) were injected bilaterally in the flanks of the nude mice. Tumor size was measured twice a week using calipers and tumor volume calculated using the formula 0.5 × (Length × width^2^). Once mice in the control group reached endpoint tumor volume of 2000mm^3^, mice were euthanized using CO_2_ and tumors were collected from all groups, weighed, and flash frozen in liquid nitrogen for downstream western blot analysis.

### Cell Titer Glo

Cell viability was measured using the CellTiter-Glo Luminescent Cell Viability Assay (Promega) according to the manufacturer’s instructions. Assays were performed at the time points indicated in the main manuscript; per well, 20 µL of reconstituted CellTiter-Glo reagent was added to 20 µL of cells in growth medium (± compound), the plate was left at room temperature for ∼10 min to stabilize the signal, and luminescence was read on a Tecan Spark microplate reader. Where indicated, raw signals were background-subtracted and normalized to vehicle controls.

### Syngeneic Orthotopic PDAC Mouse Model

Mouse studies were conducted at the animal facility of Sanford Burnham Prebys Medical Discovery Institute (SBP) in accordance with the mouse handling and experimental protocols that were approved by SBP Institutional Animal Care and Use Committee (IACUC, approval number 20-073). Six-week-old female C57BL/6 mice were purchased from Jackson Laboratories. The syngeneic orthotopic PDAC mouse models were established as previously described ^54^. Briefly, 2.5 x 10^4^ KPC.4662 cells were suspended in PBS with 50% matrigel (Corning) and injected into the pancreata of C57BL/6 mice. Ten days post inoculation, mice were randomly grouped and administrated with vehicle or 40 mg/kg **SBP-1750** by oral gavage five days a week. On day 12 and day 27, blood was collected. On day 28, mice were euthanized. Tumors were harvested, and tumor weights were measured. Tumor tissues were collected for PD studies and immunohistochemistry (IHC) analysis. Tumor macrometastases in the liver and intestine were visually inspected and recorded.

For the PD studies, tissue samples were frozen in liquid nitrogen and in vivo target engagement was assessed by immunoblotting. Total protein extracts were prepared in T-PER™ Tissue Protein Extraction Reagent (Thermo Scientific, #78510) containing a protease inhibitor cocktail (Thermo Scientific #A32953). The lysates were immunoblotted with Atg13 (D4P1K) (CST #13273) and Atg101 (E1Z4W) (CST #13492) antibodies. GAPDH (14C10) antibody (CST #2118) was used as loading control. Images were obtained on a ChemiDoc Imaging System (Bio-Rad) and Bio-Rad Image Lab software 6.1 was used for densitometry analyses with background subtraction at 50.0 mm (disk size) and band detection sensitivity at 50%. Atg13, Atg101, and GAPDH protein signal densities (i.e., adjusted total band volume) were quantified for vehicle control and experimental samples.

IHC staining was performed as previously described ^54^. Formalin-fixed paraffin-embedded (FFPE) tumor tissues were cut to 5 μm sections and heated in 0.01 M citrate buffer (pH 6.0) for heat-induced antigen retrieval. For CD4 and CD8 staining, tumor sections were blocked with 10% goat serum plus 1% BSA at room temperature (R.T.) for 1 hour and then stained with anti-CD4

(Abcam, 1:500) or anti-CD8 (Abcam, 1:1500) antibodies at 4 °C overnight, followed by incubation with biotinylated goat anti-rabbit secondary antibody (Vector Labs, BA-1000, BA-9400) at R.T. for 1.5 hours. The staining signal was developed using HRP-coupled ABC reagent (Vector Labs) and the DAB HRP Substrate Kit (Vector Labs).

### PK Methods

All experimental protocols were approved by the Institutional Animal Care and Use Committee (IACUC) of Sanford Burnham Prebys Medical Discovery Institute and performed in conformity with regulations and guidelines of the National Institutes of Health (NIH) and the Association for Assessment and Accreditation of Laboratory Animal Care (AAALAC). Eight-week-old female C57BL/6J mice were housed in ventilated cages with temperature ∼25°C, 40%-70% humidity, and a 12-h light/12-h dark cycle. The mice were fed a standard laboratory diet and food and water were supplied ad libitum. Mice were dosed with SBP-1750 in 5% DMSO, 10% Tween 80, and 85% deionized H_2_O via oral gavage (PO, 10 mg/kg). At 15-30 min and 1, 2, 4, 8, and 24 h post dosing, blood samples were collected retro-orbitally, and plasma was separated by centrifugation. Plasma samples were extracted with acetonitrile: water (4:1) with 0.1% formic acid containing verapamil as an internal standard. Compound concentration was determined by LC-MS/MS on a Shimadzu Nexera X2 HPLC coupled to an AB Sciex 6500 QTRAP, and data was analyzed using PKSolver software.

### Cas9, gRNA, and donor DNA for Creation of A549 ATG13 HiBiT

Alt-R S.p. Cas9 Nuclease V3, Alt-R CRISPR RNA (crRNA), Alt-R transactivating crRNA (tracrRNA), Nuclease-Free Duplex Buffer, and HDR DNA Oligos were obtained from Integrated DNA Technologies (IDT). 3.6 µL of Cas9 was diluted from its 62 µM stock in 2.4 µL Neon Electroporation Resuspension Buffer R per reaction. gRNA was prepared by combining 1.65 µL of 200 nmol crRNA, 1.65 µL of 200 nmol tracrRNA, and 4.2 µL Nuclease-Free Duplex buffer for a final concentration of 44 µM per reaction. This was then incubated at 95 °C for 5 min and then cooled to room temperature. dsDNA HDR donor templates were resuspended in nuclease-free IDTE Buffer, + and – strands were combined in equal volume and incubated at 95 °C for 5 min and then cooling to room temperature.

### Generation of HiBiT knock-in pools

RNP complexes were assembled by incubating 6 µL (264 pmol) Cas9 and 6 µL (216 pmol) gRNA for 10 min at ambient temperature. 5 × 10^6^ A549 cells were resuspended in 90 µl of Neon Electroporation Resuspension Buffer R. 10 µL of RNP complex, 1µL 100 pmol of donor DNA, and 2 µL of 10.8 µM Alt R Electroporation Enhancer were added to the cell suspension. Cells were electroporated with the Neon Transfection System (Thermo Fisher Scientific) using 100-µL tips (1,200 V, two 30-ms pulses) and immediately plated into six-well dishes in antibiotic-free growth medium (2 mL per well). After 48 h, cultures were returned to routine medium and single-cell clones were established by cell sorting into 96-well plates. Clones were expanded and replica-plated; HiBiT-positive populations were identified with the HiBiT Lytic Detection System (Promega) and correct on-target tagging was verified by immunoblotting with the HiBiT-specific antibody provided in the HiBiT Blotting System (Promega, N2410)

### In Cell Western Blots

A549 cells (2,000 per well) were seeded in black, clear-bottom 384-well plates (20 µL RPMI per well). After 24 h, cells were treated by adding 20 µL of 2× compound stocks prepared in RPMI. Following 48 h at 37 °C, plates were washed on an ELx washer, overlaid with PBS (20 µL), and fixed for 20 min in formaldehyde (final 1.85%). Fixed cells were washed with PBS, permeabilized in 0.1% Triton X-100 (three 10-min washes), and blocked with Odyssey Blocking Buffer (20 µL per well, 1 h, room temperature). Primary antibodies diluted in Odyssey Blocking Buffer were applied overnight at 4 °C: anti-ATG13 (E1Y9V, Cell Signaling Technology #13468; 1:250) and anti-α-tubulin (CST #3873; 1:750); blank wells received buffer (Odyssey Blocking Buffer:PBS, 1:1). After three 10-min washes in 0.1% Tween-20, secondary antibodies (LI-COR IRDye 680RD goat anti-rabbit and IRDye 800CW goat anti-mouse; each 1:800 in blocking buffer) were incubated for 1 h at room temperature in the dark. Plates were washed (three times, 0.1% Tween-20) and imaged on an Odyssey imager (LI-COR)

### Lytic HiBiT detection

A549 ATG13 HiBiT cells were plated at confluency in white clear bottom tissue culture plates (Corning #3917) in 20 µL (384 well assay) or 100 µL (96 well assay) growth medium. The day after seeding, equal volume of 2x compound was added to the cells and allowed to incubate at 37 C for determined amount of time. HiBiT was detected using the Nano-Glo HiBiT Lytic Detection System (Promega N3030) following manufacturer’s instructions. After compound incubation, plates were washed with PBS and media + compound is replaced with 20 µL (384 well) or 50 µL (96 well) of PBS. 20µL (384 well) or 50 µL (96 well) of HiBiT Lytic Buffer with 1:50 Lysis Buffer and 1:100 LgBit protein were added to experimental wells. Background wells were measured in buffer without LgBit protein. Cells were shaken on an orbital shaker for 3 mins and then left to incubate for 30 mins before recording luminesce on a Tecan Spark with 1.5 s integration time.

### Cell Titer Fluor Assays

5,000 cells per well were plated in black clear bottom tissue culture plates (Corning #3542) in 20 µL (384 well assay) growth medium. The day after seeding, equal volume of 2x compound was added to the cells and allowed to incubate at 37 C for 6 hours. HiBiT was detected using the Cell Titer Fluor Cell Viability assay (Promega G6080) following manufacturer’s instructions. After compound incubation, plates were washed with PBS and media + compound is replaced with 20 µL (384 well), after which equal volume of Cell Titer Fluor reagent is added to each well. Cells were shaken on an orbital shaker for 3 mins and then left to incubate at 37 C for 30 mins before recording fluorescence on a Tecan Spark.

### Z’ Factor Calculation

The Z’ factor was calculated to evaluate the assay’s suitability for high-throughput screening in both 384-well and 1536-well formats. For 384-well plates, untreated wells represented the maximum signal, while wells containing only lytic reagent without cells served as the minimum signal control. For the pilot screen in 1536-well plates, 3 µL of cells and 3 µL of HiBiT detection reagent were used per well. Luminescence readings were obtained using a Tecan Spark plate reader, and Z’ factor values were derived from the signal variability and separation between controls. The assay consistently yielded Z’ values ≥ 0.5, demonstrating its robustness for screening.

The Z’ factor is a statistical measure of assay quality, calculated using the standard deviation and mean values of both positive and negative controls. A Z’ factor between 0.5 and 1.0 indicates an excellent assay with high signal separation and minimal variability, making it ideal for high-throughput applications. A Z’ factor of 0.71 signifies that our assay is highly reliable and reproducible, with sufficient dynamic range to distinguish between varying degrees of ATG13 degradation in response to different compounds. In addition, we have successfully adapted the ATG13 HiBiT assay for use in a 1536-well format, further increasing its potential for ultra-high-throughput screening (**Supplemental Table 2**).

### GFP-LC3-RFP-LC3ΔG Autophagic Flux Assay

A549 cells were transfected with the GFP-LC3-RFP-LC3ΔG plasmid (Addgene plasmid #84572) and treated with compounds at 5 µM in EBSS starvation media for 18 hours. Fluorescence was measured using the BD LSRFortessa Flow Cytometer, capturing the percentage of RFP positive cells which were also GFP positive for each condition. Data were normalized to untreated cells in normal growth media (0% autophagic flux) and cells in EBSS without compound (100% autophagic flux). Data was generated to assess the relationship between compound concentration and autophagy inhibition, correlating fluorescence changes to autophagic flux.

### Cell Viability Assay Using CellTiter-Glo®

Cell viability was assessed using the CellTiter-Glo® Luminescent Cell Viability Assay (Promega, G7570), which quantifies ATP levels as a proxy for metabolically active cells. Compounds were pre-plated in black-walled, clear-bottom 384-well plates (Corning #3764). A549 cells were then seeded directly onto the plates at 20 µL per well. Plates were incubated at 37°C under standard growth conditions for 24 or 48 hours. After incubation, 20 µL of CellTiter-Glo® reagent was added to each well. The plates were shaken gently on an orbital shaker for 2 minutes to ensure cell lysis and then incubated at room temperature for 10 minutes to stabilize the luminescent signal. Luminescence was measured using a Tecan Spark plate reader with a 1-second integration time per well. Data were analyzed to generate dose-response curves, and IC50 values were determined using non-linear regression analysis in GraphPad Prism.

## SUPPORTING INFORMATION

A Figure showing the structures of SBI-0206965, SBP-7455, SBP-7501, and SBP-5147, a Scheme showing the general synthetic route for the preparation of ULK1/2 inhibitors, a Figure showing the validation of ULK knockdown, a Table showing oligonucleotide sequences utilized for CRISPR knockout of ULK1 and ULK2, and a Table showing the Z’ Factor Analysis for the 1536 Well Plate A549 ATG13 HiBiT Assay.

## AUTHOR INFORMATION

Corresponding Author, **

Author Contributions

The manuscript was written through contributions of all authors. All authors have given approval to the final version of the manuscript.

Notes

The authors declare no competing financial interest.

## ACKNOWLEDGMENTS

This work was supported by The Epstein Family Foundation (N.D.P.C.), Pancreatic Action Network Foundation Grant 19-65-COSF (N.D.P.C. and R.J.S), 2022 Curebound Discovery Grant (N.D.P.C. and R.J.S), and 2023 CureBound Discovery Grant 23TG05 (N.D.P.C. and R.J.S). Research reported in this publication was supported by the National Cancer Institute of the National Institutes of Health under Award Number P30CA030199. The authors would like to thank the staff of the SBP Shared Resources Flow Cytometry Core and Animal Facility for their support in experimental design and analyses under this Cancer Center Support Grant. The content herein is solely the responsibility of the authors and does not necessarily represent the official views of the National Institutes of Health. This study was also supported by grants to R.J.S. from the National Institute of Health (R35CA220538 and P01CA120964). S.N.B. was supported by NIH postdoctoral fellowship (F32CA206400).

## SUPPLEMENTAL DATA

**Supplemental Figure 1:**
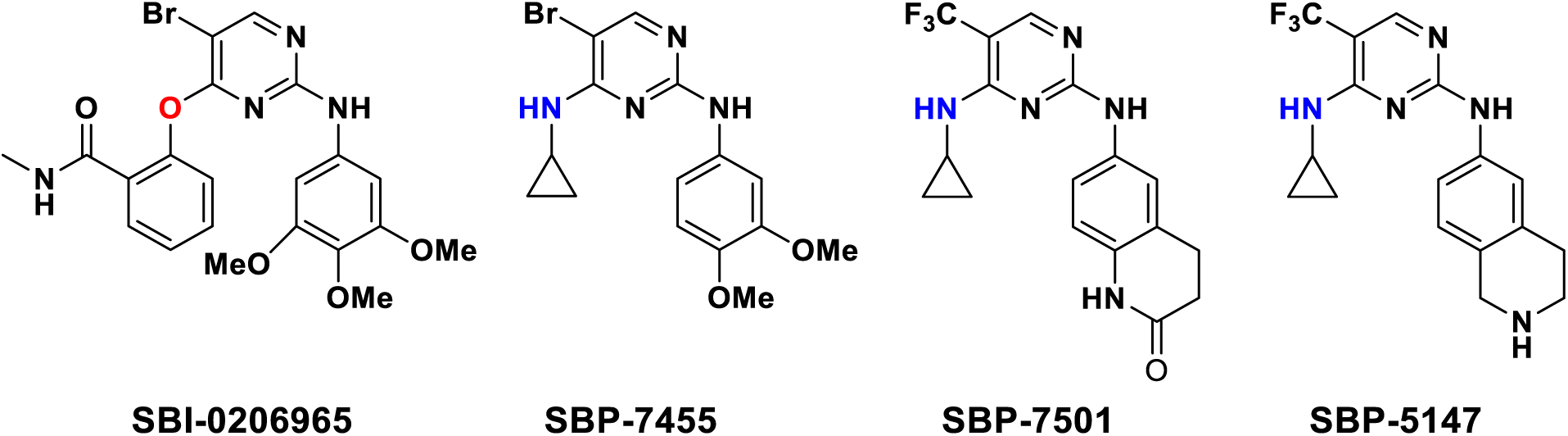
Structures of SBI-0206965, SBP-7455, SBP-7501, and SBP-5147.

**Supplemental Scheme 1:**
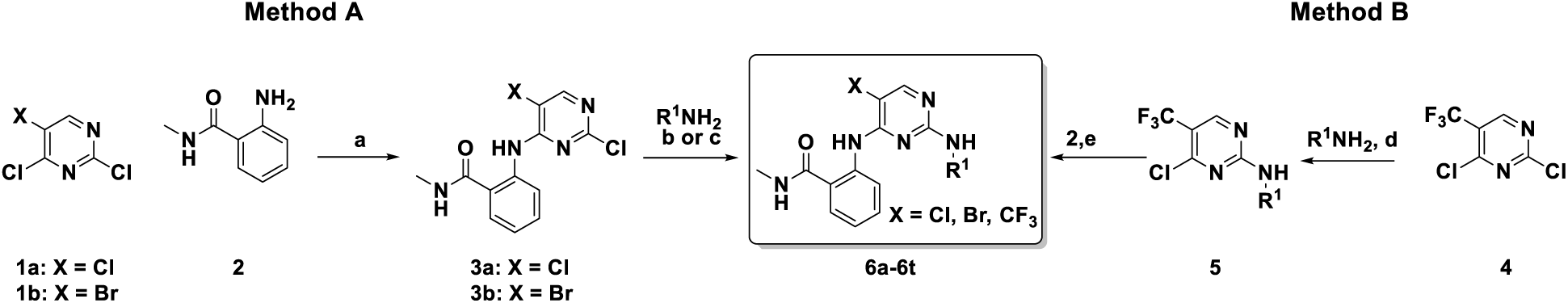
General synthetic route for the preparation of ULK1/2 inhibitors. Reagents and conditions: (a) K_2_CO_3_, DMF, 75 °C, 5 h (b) ^n^BuOH, 110 °C, 16-24 h (c) HCl, EtOH, 60 °C, 6 h. (d) ZnCl_2_ (1.0 M solution in ether), CH_2_Cl_2_-*^t^*BuOH, 0 °C, 1 h; R^1^NH_2_, Et_3_N, 0 °C-rt, 1.5 h (e) ^n^BuOH, DIEA, 110 °C, 16-24 h.

**Supplemental Figure 2:**
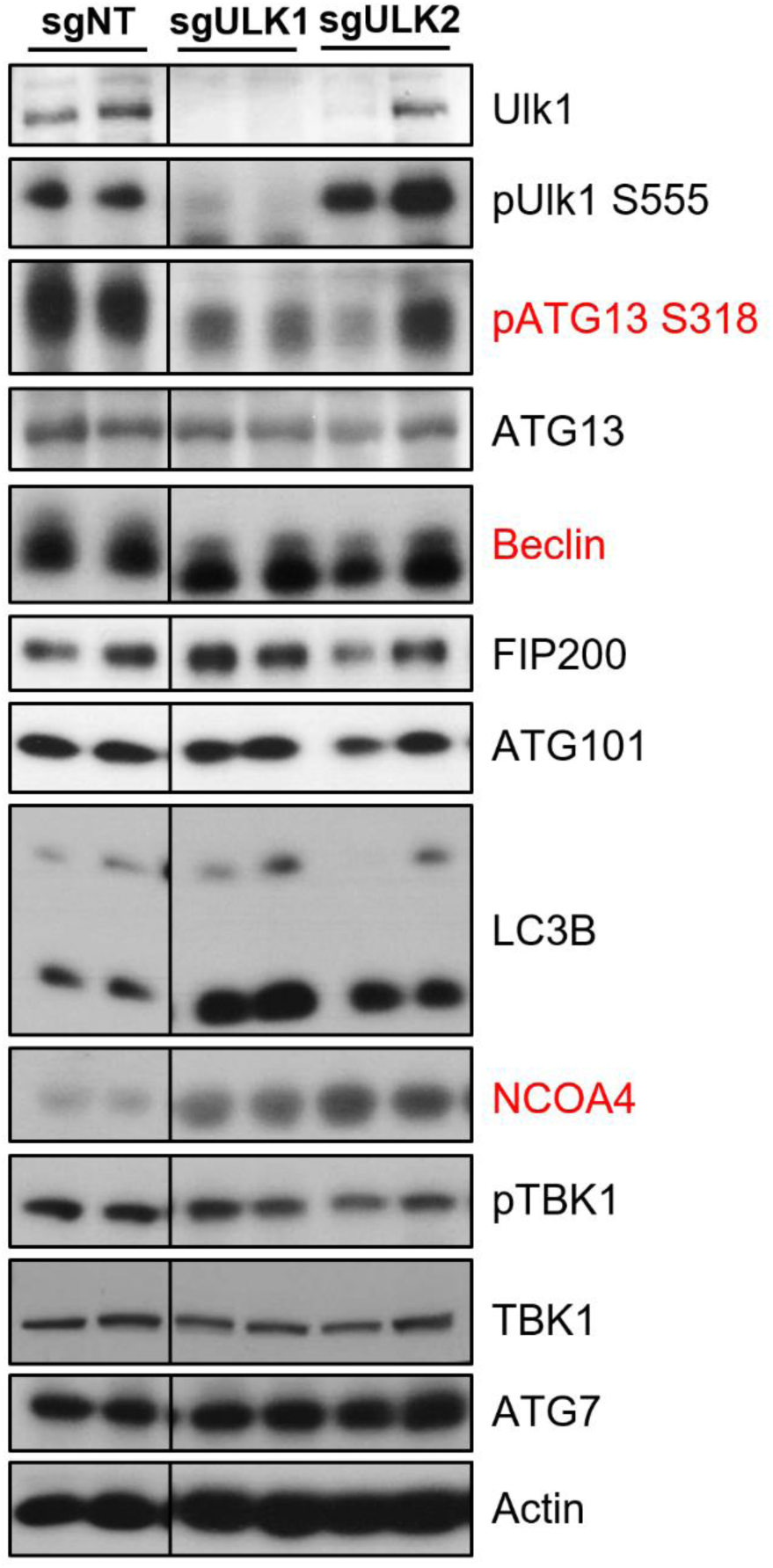
Validation of ULK Knockdown. Western blots on lysates from MiaPaca2 cells stably expressing small guide RNAs (sg) targeting ULK1 or ULK2 alone, or lines targeted to co-delete ULK1 and ULK2 demonstrate efficient deletion. Direct ULK1 substrates are highlighted in red. Control sg-RNA treated MiaPaca2 cells (sgNT) as well as lines expressing guides targeting critical autophagy regulators ATG5, ATG7, and FIP200 are also shown for comparison.

**Supplemental Table 1.**
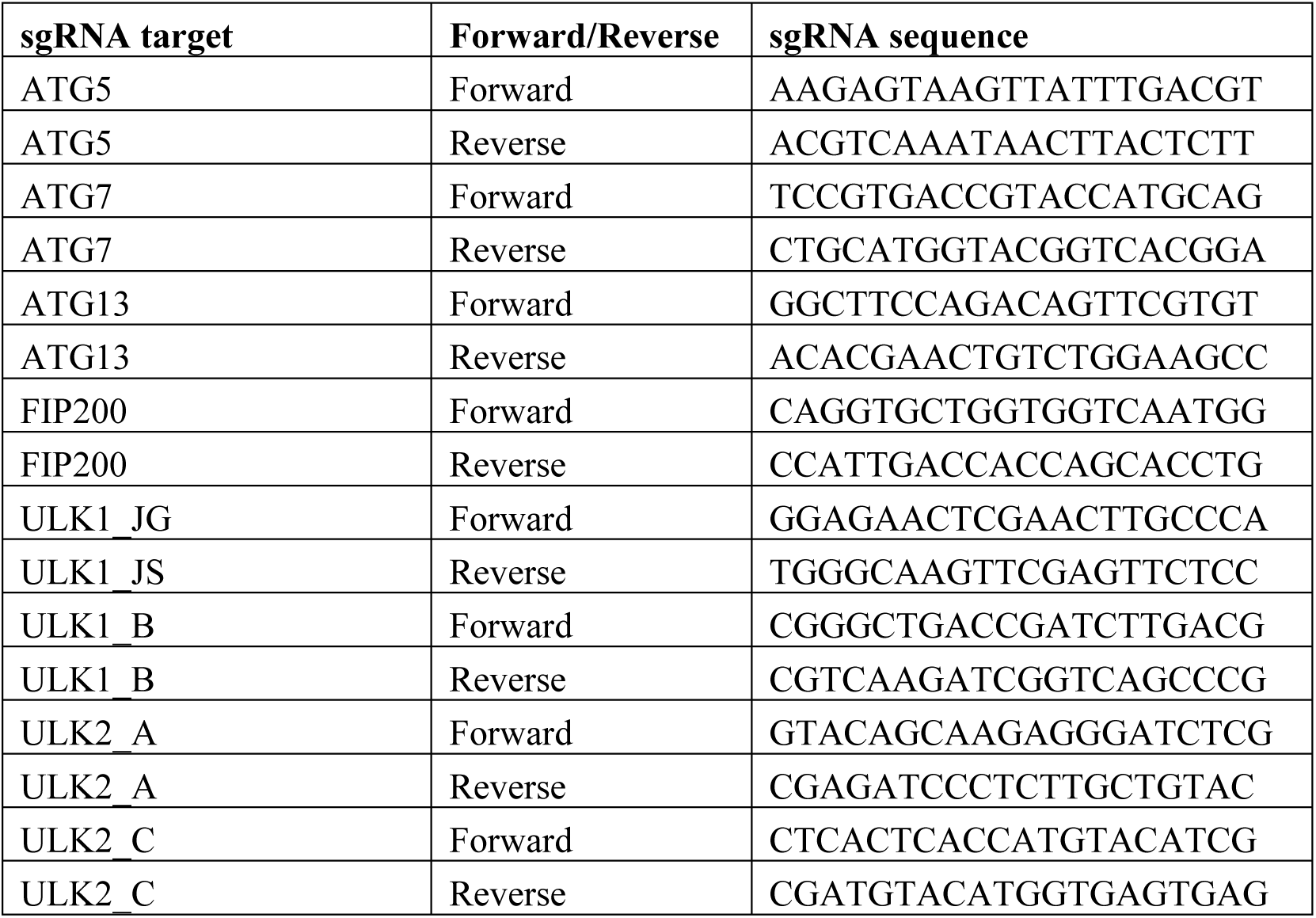
sgRNA Targeting Sequence for Autophagy Genes.

**Supplemental Table 2:**
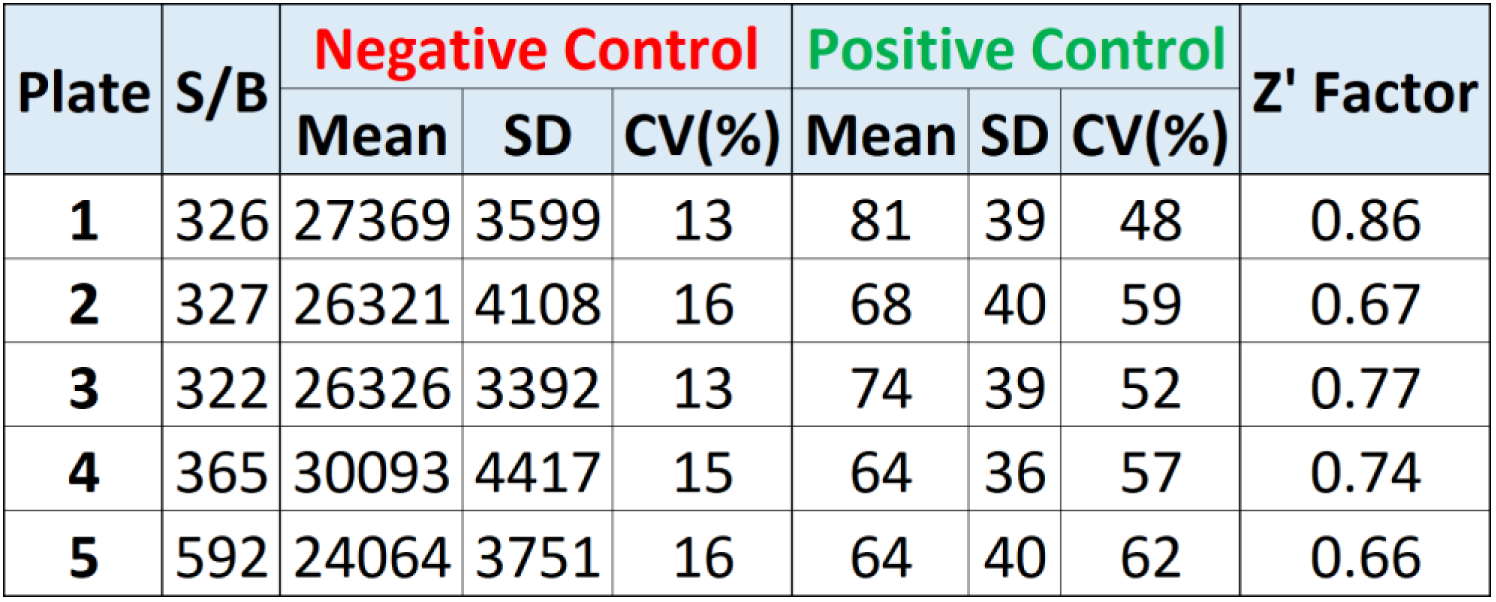
Z’ Factor Analysis for the 1536 Well Plate A549 ATG13 HiBiT Assay. A549 ATG13 HiBiT assay optimized to the 1536 well format. Pilot screen z’ factor average = 0.74. 7.5 nL novel compounds, 3 nL A549 ATG13 HiBiT Cells, 6 h incubation at 37 C, 3 nL of HiBiT lytic reagent.

